# Modular Surface Engineering of Feline Parvovirus Virus-Like Particles for Antigen Presentation and Receptor-Mediated Cell Targeting

**DOI:** 10.64898/2026.09.09.750088

**Authors:** Antonina Naskalska, Artur P. Biela, Jan Różycki, Kinga Fic, Edyta Kuś, Jakub Nowak

## Abstract

Virus-like particles (VLPs) are non-infectious, multiprotein nanostructures that mimic the architecture of viruses while lacking genetic material. Their structural versatility and safety profile make them promising platforms for biomedical applications, including vaccine development, targeted drug delivery, and diagnostics. Among various functionalization strategies, click-chemistry has emerged as a precise and efficient method for attaching bioactive molecules to VLP surfaces. In this study, we report the production and functionalization of VLPs derived from the feline panleukopenia virus (FPV), a parvovirus closely related to the well-characterized canine parvovirus (CPV). We demonstrate successful expression, assembly, and purification of FPV VLPs, followed by their modification with fluorophores, peptides, oligonucleotides, and proteins using copper-free click-chemistry. Furthermore, using cryo-EM, we structurally characterize FPV VLP functionalized with antigenic peptides, providing detailed insights for further development of such vaccine prototype. Finally, we demonstrate that our FPV VLP specifically transduce feline and human cells expressing transferrin receptor type 1 (TfR), thus validating their potential as a unique platform for future nanotechnological and therapeutic applications.

## 1. Introduction

Virus-like particles (VLPs) are multiprotein structures that resemble native virus particles in morphology and conformation. However, because they lack viral genetic material, they are non-infectious and safe to handle. At the same time, their composition of viral structural proteins enables them to retain the immunogenicity and receptor specificity of their parental viruses. In addition, VLPs are highly amenable to functionalization with a variety of molecular entities, rendering them versatile platforms for the development of customized nanocarriers for numerous biomedical and nanotechnological applications [1, 2]. Namely, functionalized VLPs are being developed as vaccines, as vehicles for targeted drug delivery, and as diagnostic tools [3–7]. Among several methods enabling functionalization of VLPs, the click-chemistry is emerging as the most robust and versatile. The advantage of click-chemistry approach is the possibility of the precise attachment of a range of bioactive molecules, not limited to proteins and peptides, as in case of genetic fusions. Examples of molecules conjugated to VLP’s surface by click-chemistry include peptides [8–11], oligonucleotides [12, 13], antibodies [14], and other proteins [15–17]. Another advantage of click-chemistry is the possibility of carrying out the reaction in commonly used aqueous buffers instead of organic solvents (often required for traditional conjugation methods), thereby significantly reducing the risk of altering the structural properties of the VLP or the functionalizing molecule. Furthermore, recent advancement in this technology eliminated the need of using copper, which might have been a limitation for biological applications [18].

Here we show production of a VLP built of capsid proteins from feline panleukopenia virus (FPV) and its modular functionalization by amine-ester coupling combined with click-chemistry.

FPV is a member of the *Parvoviridae* family and is closely related to the first isolated and characterized parvovirus: canine parvovirus (CPV) [19, 20]. FPV causes severe gastrointestinal disease and immunosuppression in infected cats, followed by high mortality. Fortunately, efficacious vaccines are available, offering long lasting protection of vaccinated animals [21]. Similarly to CPV, FPV infects cells through binding the transferrin receptor type 1 (TfR) and this interaction determines virus host range [22, 23]. Interestingly, both CPV and FPV retain the ability to infect cells expressing human TfR in laboratory conditions [24], although no real infection of a human by CPV or FPV has been ever reported.

The structure of CPV is well studied, providing insights in the capsid composition, which is built of 60 copies of two main proteins: VP2 (more abundant) and VP1 [25–28]. Importantly, FPV which is phylogenetically parental (and similar) to CPV, shares genetic and structural similarities with CPV [29, 30]. Observation that VLPs (called *empty capsids* at that time) are formed together with infectious viral particles during cell culture infection (with CPV or FPV) has been reported as early as in 1993 [31–33] and provided fundamental structural basis for further development of these particles. Notably, Parrish and colleagues noticed that VP2, being in fact truncated VP1, is sufficient to form VLPs. The potential utility of CPV VLPs as a displaying platform for heterologous antigens has been initially explored by researchers from a biotech company Ingenasa [34–36] and subsequently expanded by other groups [37–40], with one notable example of exploring CPV VLPs as tumor targeting vector [41, 42]. Recently, recombinant expression of FPV VLPs and their immunogenic assessment as vaccine candidates has been reported [43–45].

Overall, the plasticity of CPV VLP and FPV VLP in terms of modifications, together with their unique ability to interact with human TfR make them attractive candidates for further development for nanomedicine purposes. In this work, we produced recombinant FPV VLPs; we extensively assessed their stability and - by the means of click-chemistry – we decorated these particles with a range of molecules: heterologous proteins; peptides; oligodeoxynucleotides; and fluorophores. Using several approaches, we estimated the efficiency of such functionalization. Furthermore, we confirmed that our FPV VLPs specifically transduce human cells expressing TfR.

## 2. Results

### 2.1. Rational design, production and purification of FPV VLPs

When designing the building blocks of FPV VLP we based on the known structure of FPV and CPV empty capsids [**Supplementary Figure 1A**] and published work on development on CPV VLP, providing evidence that up to 240 amino acids might be fused to the N-terminus of VP2 protein, without interfering capsid assembly, and resulting in external exposure of the appended tag/peptide [37–39, 46, 47]. Since sequences of the VP2 from FPV is highly similar to that from CPV, it is probable that FPV-derived VLP would tolerate analogous capsid modifications to CPV-derived VLPs. Therefore we fused the His-tag to the N-terminus of FPV VP2 protein, expecting that such a design will allow for affinity purification of the assembled VLP. At the same time we also prepared a VP2 version without the His-tag (in the case it interferes with VLP assembly) [**Supplementary Figure 1B**].

Both versions of VP2 protein were expressed in insect cells, resulting in the proper assembly of particles, as demonstrated by further characterization of the obtained samples [**Figure 1A**]. Namely, their fractionation by size exclusion chromatography (SEC) gave a peak at the expected elution volume (EV), for the used Superose 6 column (EV = 12 mL, corresponding to molecular weight in the range of MDa) [**Figure 1B**]. Proteins from the peak fractions appeared to migrate in native PAGE with the similar pattern to our previously characterized VLPs and protein cages [48, 49], indicating presence of larger assemblies [**Figure 1C**]. In the case of FPV VLPs composed of His-VP2 proteins, SEC was preceded with sample purification on Ni^2+^ resin. Fractions from this column were subjected to Western blot analysis and detection with anti-His antibodies. Observed signal from the band migrating at ~ 70 kDa corresponds to the molecular weight of FPV His-VP2, being 67.7 kDa [**Figure 1D**]. Finally, purified samples were imaged using transmission electron microscopy (TEM), providing eventual evidence that the obtained material are properly assembled VLPs [**Figure 1E**].

**Figure 1.**
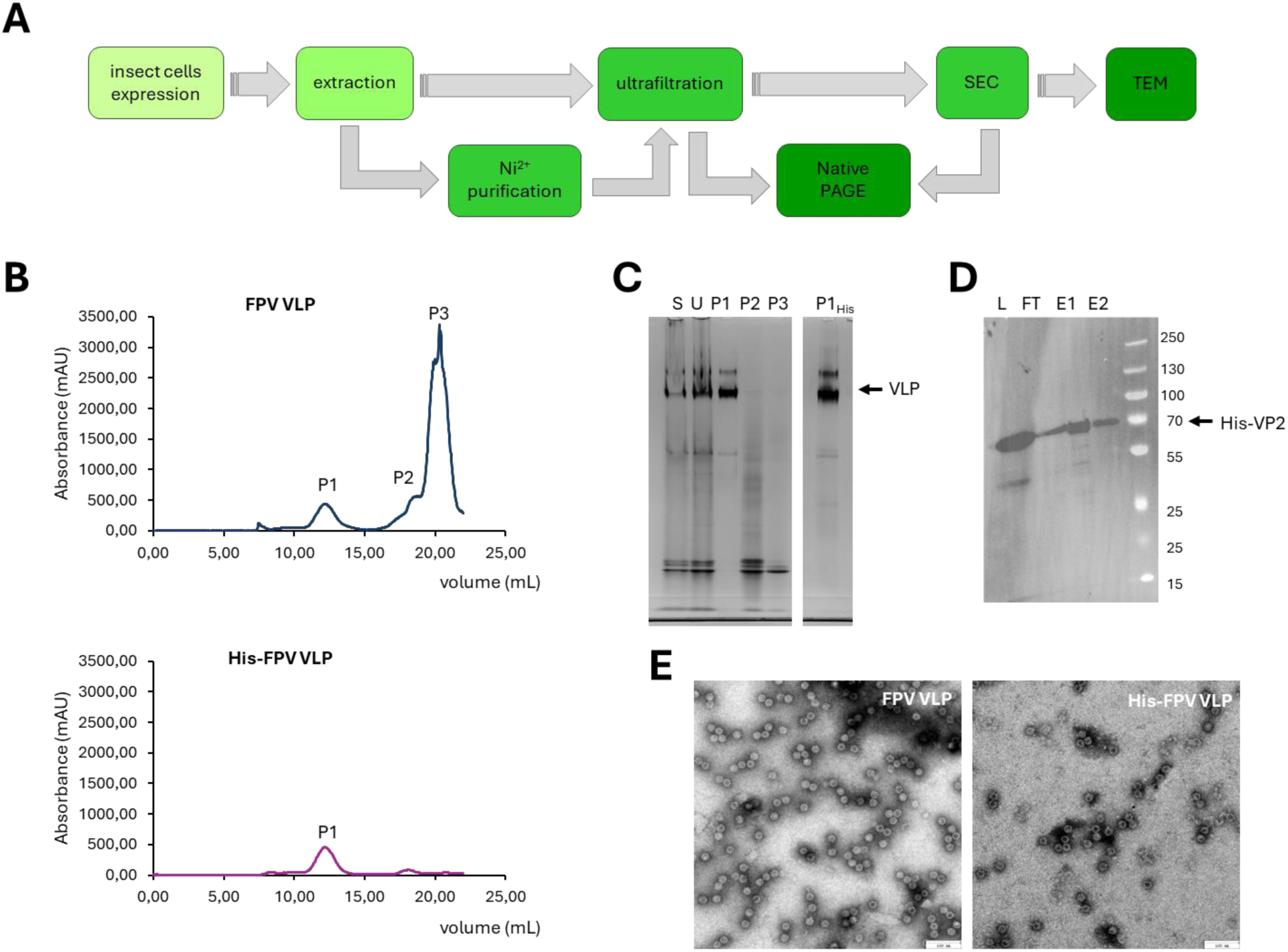
Production of FPV VLPs. Diagram showing the procedure of FPV VLP production: VP2 or His-VP2 proteins are expressed in insect cells, then extracted by sonication followed with centrifugation. Assembled VLPs present in the supernatant are separated from smaller proteins by ultrafiltration. Additional step of affinity (Ni^2+^) purification is performed in the case of His-VP2 protein. VLPs are further purified using size exclusion chromatography (SEC), subjected to native polyacrylamide gel electrophoresis (PAGE) and then inspected under transmission electron microscope (TEM) (**A**). SEC profiles of FPV (upper) and His-FPV (lower) VLPs (**B**). Native PAGE separation of supernatant after extraction (S); eluate after ultrafiltration (U); fractions from peak 1 (P1); 2 (P2); and 3 (P3) collected from SEC (**C**). Fractions collected after purification of His-VP2 protein using affinity chromatography: L – load; FT – flow through; E1, E2 – elution, separated by SDS-PAGE and detected in Western blot with anti-His antibody (**D**). TEM images of purified FPV (left) and His-FPV (right) VLPs (**E**).

### 2.2. Stability of FPV VLPs

To further characterize obtained FPV VLPs we tested their resistance to high temperatures, pH range, detergents, polyethylene glycol (PEG), denaturing agents, and organic solvents. VLPs exposed to the specific conditions were subsequently subjected to native PAGE, together with the untreated control [**Figure 2**]. As can be observed from the stained gels, FPV VLPs (both His-tagged and untagged variant) tolerate up to 60°C, at least for 1 hour [**Figure 2A**]. They also tolerate strongly acidic environment (pH 1) and remain intact when alkalized up to pH 9.5 [**Figure 2B**]. Addition of Tween 20 or Tween 80 (up to 0.2 % w/V) detergents, as well as PEG 8000 (up to 8 % w/V) has no effect on FPV VLPs integrity, judging from their migration pattern in native electrophoresis [**Figure 2C**].

**Figure 2.**
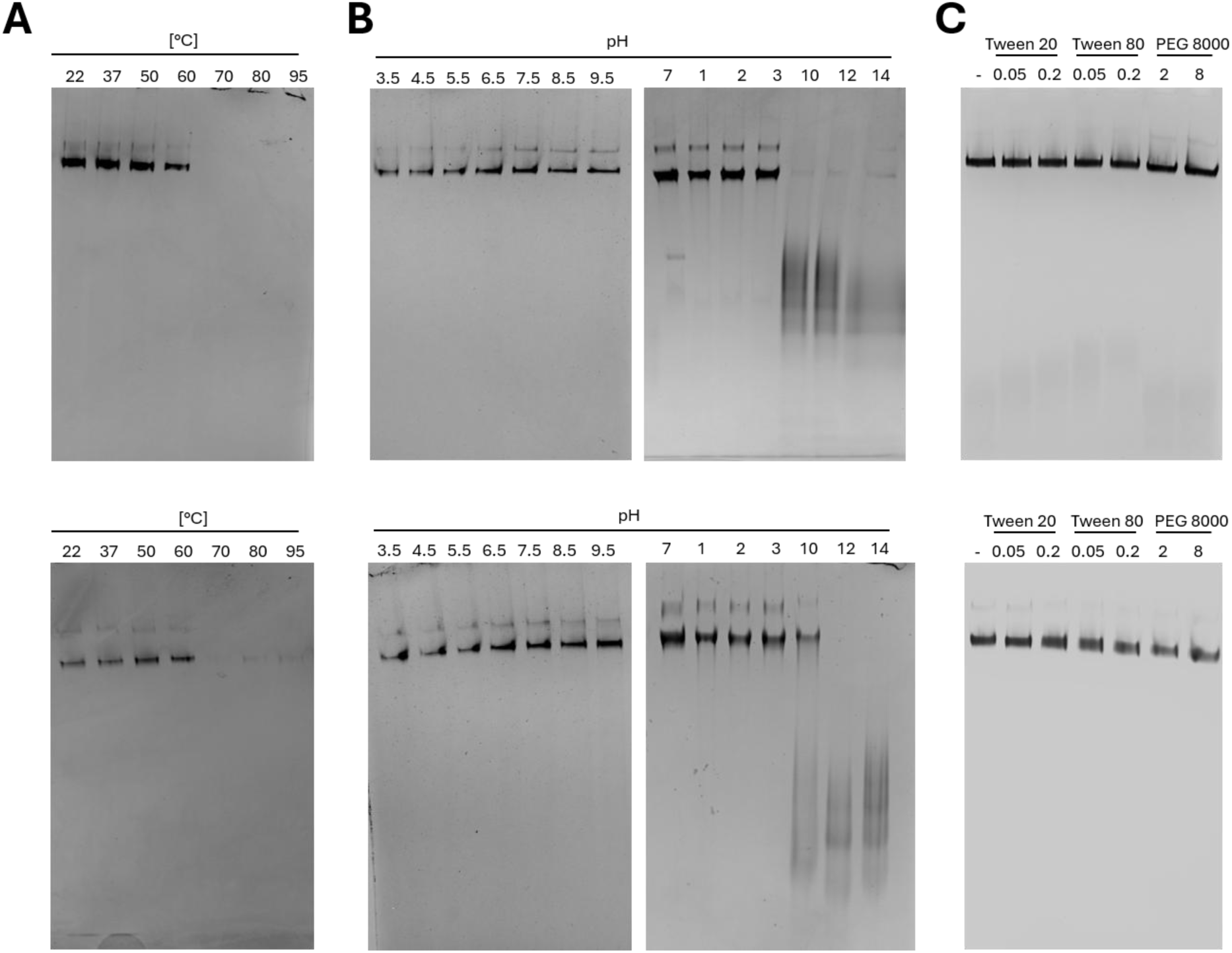
Stability of FPV VLPs in temperature, pH range, and presence of detergents. Stability of FPV VLPs (upper) and His-FPV VLPs (lower) in high temperatures (**A**); a range of pH (**B**); addition of Tween 20, Tween 80 and PEG 8000 (**C**) was assessed by native PAGE, as indicated.

Anticipating the need to use organic solvents and denaturing agents in subsequent analyses, we also examined their effect of FPV VLPs. As shown in **Supplementary Figure 2** FPV VLPs seem unaffected by the presence of DMSO and DMF (up to 25% V/V), and resist in SDS (up to 4% w/V), 1 M urea, guanidine chloride (up to 2.5 M). Interestingly, FPV VLPs and His-FPV VLPs exhibit some differences in tolerance to denaturing agents and organic solvents, namely: FPV VLPs seem to be more resistant to urea, whereas His-FPV VLPs support higher concentration of DMSO [**Supplementary Figure 2**].

To gain a deeper insight into stability of FPV VLPs in different buffer conditions we also analyzed their fluorometric transitions during thermophoresis upon incubation in buffers of the pH ranging from 4 to 9.5. As demonstrated by differential scanning fluorimetry (DSF) analysis, FPV VLPs (both with and without His-tag) support higher temperatures in acidic buffers (sodium citrate pH 4 and MES pH 5.5) than in alkaline buffers (TRIS pH 8.5 – 9.5) [**Figure 3A**], which is in accordance with native PAGE results [Figure. 2A]. Dynamic light scattering (DLS) measurement showed that mean FPV VLP and His-FPV VLP hydrodynamic radius of 14.23 nm and 16.63 nm (which correspond to 28.46 and 33.26 nm of diameter), respectively [**Figure 3B**]. While presence of the His-tag has no significant influence on VLP thermal stability, although it influences its hydrodynamic diameter (radius), possibly reflecting the impact of the His tag on overall particle hydrophilicity. Altogether, these results indicate that buffers within the range of pH 4 - 5.5 confer thermal stability and might be more suitable for VLPs storage, while pH > 6.5 improves colloidal stability which might be beneficial for downstream VLPs applications (such as conjugation reactions).

**Figure 3.**
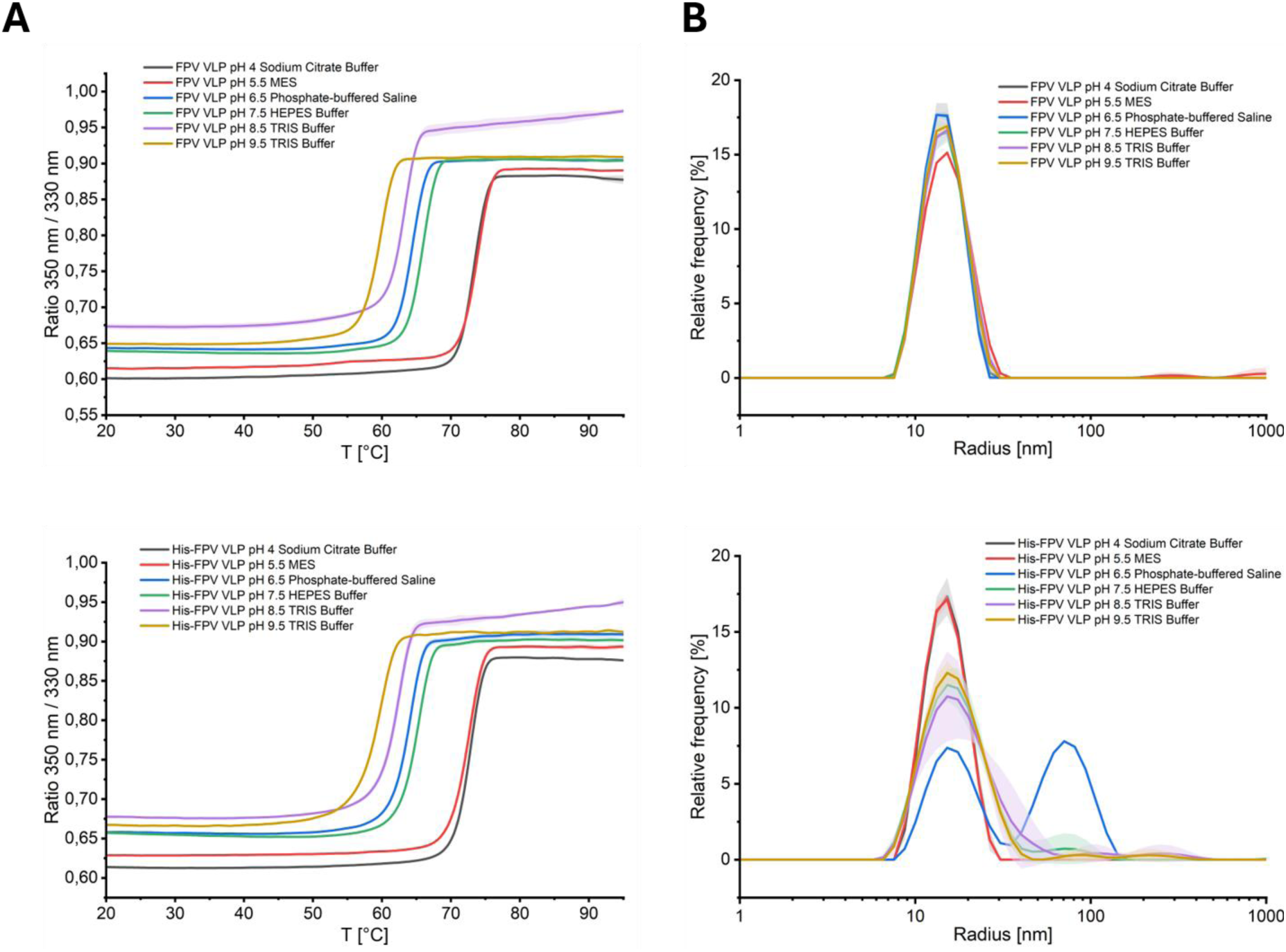
Buffer dependent thermal stability and size of FPV VLPs. Thermal unfolding profiles measured by nanoDSF, presented as intrinsic tryptophan fluorescence ratio (350/330 nm) as a function of temperature (**A**). Dynamic light scattering (DLS) analysis showing the hydrodynamic radius light distributions and relative frequency (polydispersity) in different buffers and pH conditions (**B**). Upper panels show results for FPV VLPs and lower panels – for His-FPV VLPs.

### 2.3. Functionalization of FPV VLPs

FPV VLP being stable and relatively easy to produce in endotoxin-free (bacteria-free) conditions are potentially attractive tools for several biomedical applications. Having this in mind, we aimed to functionalize them with a range of molecules, including fluorescent dyes, oligodeoxynucleotides, peptides, and heterologous proteins. With this purpose, we first examined the possibility of using reactive groups of lysines and cysteines present in the FPV VP2 protein sequence [**Supplementary Figure 3A**], potentially enabling chemical conjugation mediated by NHS esters and maleimides, respectively. Using data provided by PDB model 1C8E [50] we calculated that approximately 180 - 240 lysines (out of total 1320 lysines present in the whole FPV VLP, 3 - 4 in each VP2 unit) are externally available at the FPV VLP surface. Surprisingly, direct conjugation of NHS-fluorophores to FPV VLPs was not efficient [**Supplementary Figure 3B, C**]. On a side note, presumably neither of 240 cysteines present in the whole FPV VLP is exposed on the particle exterior, thus practically eliminating this amino acid as attachment site for external VLP modifications.

On the contrary, FPV VLPs turned out to be easily modifiable through a two steps reaction, involving click-chemistry approach: first, an adaptor molecule (NHS-DBCO ester) was conjugated to lysines accessible on the VLP surface; then azide-modified molecules were “clicked” to the adaptor molecule displaying an alkyne group [**Supplementary Figure 3C** and **Figure 4A**]. For initial screening for optimal reaction conditions we used azide-modified Cy3 fluorophore. To investigate the maximum amount of this molecule that can be conjugated to FPV VLPs, we tested different molar ratios of the reagents involved. As demonstrated by native-PAGE with fluorescent detection the reaction reaches saturation in conditions: 2 × molar excess of NHS-DBCO ester and 2 × molar excess of azide-Cy3 over accessible lysines [**Supplementary Figure 3C**]. Using the above strategy we managed to decorate FPV VLPs with oligodeoxynucleotides conjugated to Cy5 fluorophore (ODN-Cy5), peptides conjugated to fluorescein (peptide-FITC), biotin-streptavidin complex conjugated to Alexa-488 dye [**Figure 4B-D**]. The latter reaction involved in fact three steps: NHS-DBCO ester conjugation to FPV VLPs (i), followed by clicking azide-modified biotin (ii), followed by adding streptavidin labeled with Alexa-488 dye (iii) [**Figure 4E**]. Next, we verified that the excess of molecules not attached to FPV VLPs can be easily removed from the reaction mixture taking advantage of the high molecular weight of decorated particles (MDa range) as compared to low molecular weight of free dyes, peptides, ODNs or even proteins (kDa range). Buffer exchange using ultrafiltration devices [**Figure 4B-D**] or size exclusion chromatography (SEC), in the case of streptavidin-Alexa488 [**Figure 4F**], allows for efficient purification of functionalized FPV VLPs. Furthermore, during SEC separation absorbance detection at the wavelength corresponding to the excitation wavelength of the labeling dye (Alexa-488), additionally confirms efficacious FPV VLP decoration [**Figure 4F**].

**Figure 4.**
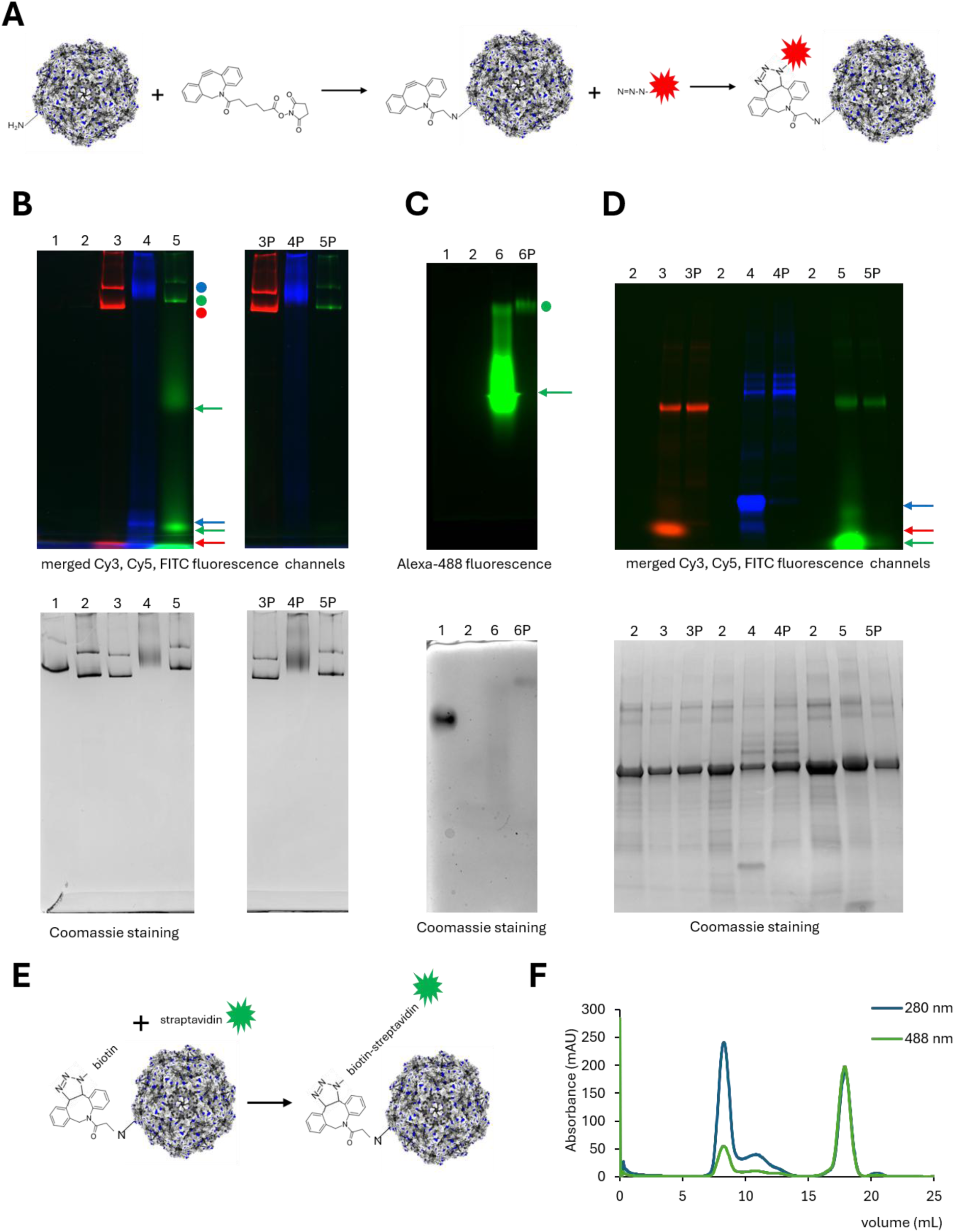
Decorating FPV VLPs. FPV VLPs functionalization was achieved by a two-step reaction: conjugating NHS-DBCO ester to lysines accessible in the VLP, followed by clicking azide-modified molecules, pre-conjugated to a fluorophore (**A**). Proper functionalisation was assessed by native PAGE (**B**), agarose gel electrophoresis (AGE) (**C**), SDS-PAGE (**D**). Gels were visualized in a fluorescent detector (upper panels) and stained with Coomassie protein dye (lower panels) to show band shift of functionalized FPV VLPs (B, C) or VP2 proteins (D). Lanes numbering (B, C, D): 1 - FPV VLP, 2 – FPV VLP-DBCO, 3 – FPV VLP-Cy3, 4 – FPV VLP-ODN-Cy5, 5 – FPV VLP-peptide-FITC, 6 – FPV VLP-biotin-streptavidin-Alexa488. Lanes marked with “P” indicate samples purified by ultrafiltration. Dots indicate decorated FPV VLPs; arrows indicate unreacted fluorescent molecules. Decoration of FPV VLPs with streptavidin involves additional step of adding streptavidin labeled with Alexa-488 dye to FPV VLP modified with biotin by prior click reaction (**E**). Size exclusion chromatography with 488 nm absorbance detection confirms FPV VLP decoration with straptavidin-Alexa-488 and allows to remove unreacted streptavidin-Alexa-488 (**F**).

Using fluorescence intensity of the signal detected in SDS-PAGE we estimated the amount of molecules present in the decorated FPV VLPs samples [**Supplementary Figure 4A**]. The calculated amount of Cy3, ODNs-Cy5, peptide-FITC molecules were: ~136, ~90 and ~39 respectively. Since this approach was not applicable for FPV VLP decorated with biotin-streptavidin-Alexa488 (due to the non-covalent nature on biotin-streptavidin interaction) we sought to quantify it by non-denaturing mass photometry (MP), however with no success. Additionally, based on HPLC analysis of FPV VLP-peptide-FITC the number of peptides loaded onto one FPV VLP was estimated to be 54 (± 5) [**Supplementary Figure 4B**]. Averaged values of decoration yields calculated using different approaches are summarized in **Supplementary Table 1**.

### 2.4. Biophysical characterization of functionalized FPV VLPs

To examine the effect of above functionalization of FPV VLPs in terms of particles size and stability, we measured their hydrodynamic radii and thermal unfolding profiles, similarly as for the unmodified FPV VLPs [**Figure 3**]. As shown in **Figure 5A** decoration with Cy3 dye; ODN-Cy5; peptide-FITC; biotin-streptavidin-Alexa-488 resulted in clear increase of particle hydrodynamic diameter by: 2 – 9 nm as compared to control FPV VLPs (with NHS-DBCO only) [**Supplementary Table 2**]. On the other hand, neither of decorations influenced particle thermal stability measured as intrinsic tryptophan fluorescence ratio (350/330 nm) as a function of temperature [**Figure 5B**]. Furthermore, inspection of decorated particles in transmission electron microscopy (TEM) confirmed their integrity [**Figure 5C**].

**Figure 5.**
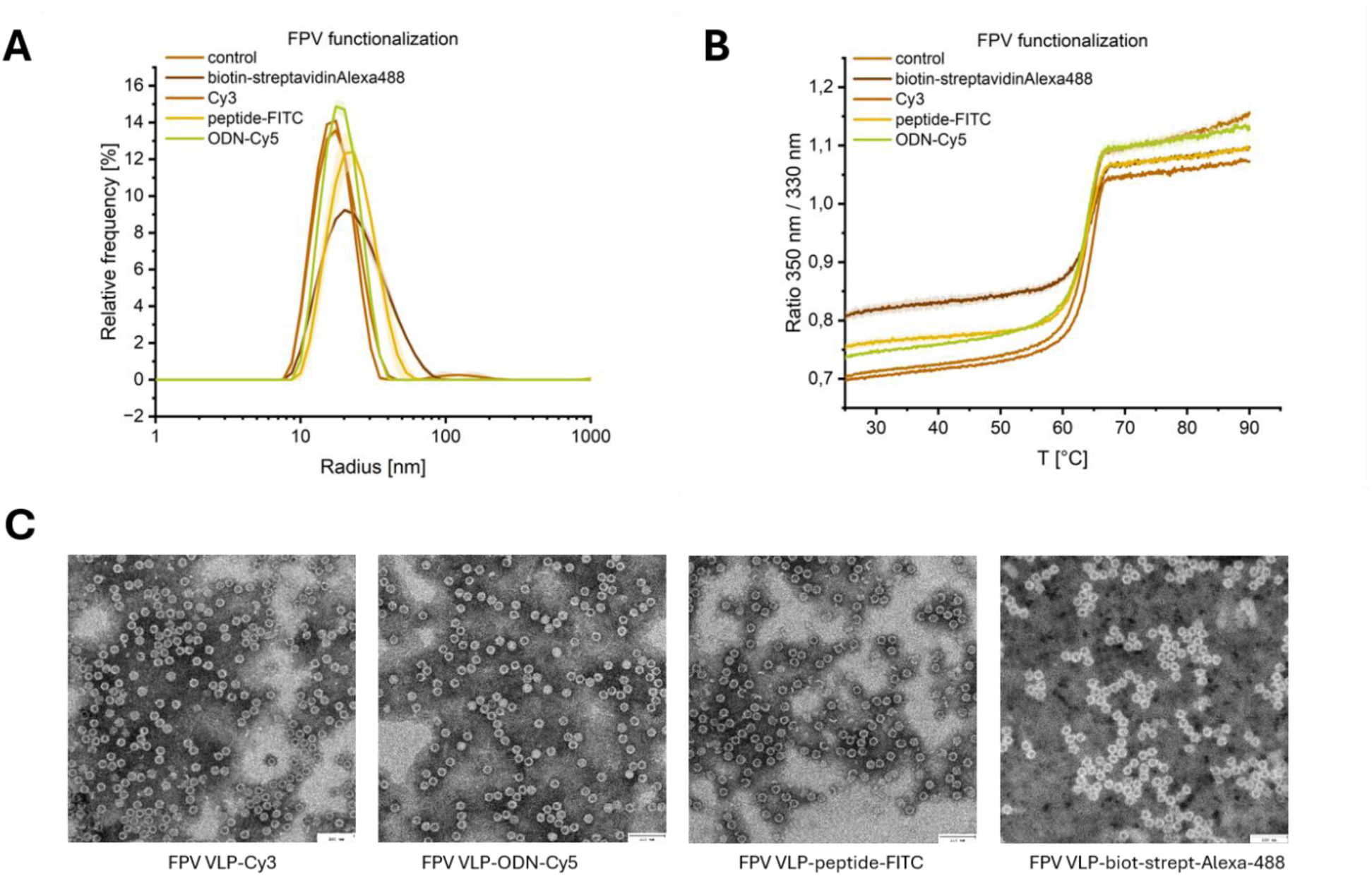
Characterization of functionalized FPV VLPs. Hydrodynamic radii of FPV VLPs decorated with Cy3 dye, ODN-Cy5 peptide-FITC, biotin-streptavidin-Alexa-488, and FPV VLPs with NHS-DBCO only were measured in DLS (**A**). Thermal stability of the same samples was assessed by nanoDSF, presented as intrinsic tryptophan fluorescence ratio (350/330 nm) as a function of temperature (**B**). Decorated FPV VLPs were inspected in TEM (**C**).

### 2.5. Structural characterization of functionalized FPV VLPs by cryo-electron microscopy

Structural characterization of FPV VLPs was performed using cryo-electron microscopy (cryo-EM). Two samples of FPV VLPs were subjected to the structural analysis: functionalized with peptide used for above (FPV VLP-peptide) and non-functionalized. To assess if the modifications introduced on the surface of the VLPs have any influence on their shape and size, and if they are visible and identifiable, head-to-head comparisons of both reconstructions were made in all applied symmetries (C1, D3, D5 and I) [**Figure 6**]. The FPV VLP without any modifications introduced on its surface (in green) does not possess any additional density on its surface, showing resolved structural features regardless of imposed symmetry to the final refinement at standard map contouring level of RMSD = 3.5. Decreasing the value of contouring level to RMSD = 1 or lower, does not show any additional densities on the surface of the reconstructed particle. In contrast, the functionalized particles (in blue) start to display additional densities (red circles) in lower (below RMDS = 1) contouring level. They are the most prominent in the reconstruction without any symmetry imposition during the refinement; they are clearly visible at RMSD = 0.75. For the other refinements with symmetry imposition, they are present at slightly lower value of RMSD; namely 0.63. This is due to the signal averaging upon the symmetry imposition. Nevertheless, in all cases (symmetries) the additional signal of the densities is present only in the case of modified FPV VLPs, proving that functionalization with a relatively small molecule (20 amino acids peptide) can be visualized by cryo-EM.

**Figure 6.**
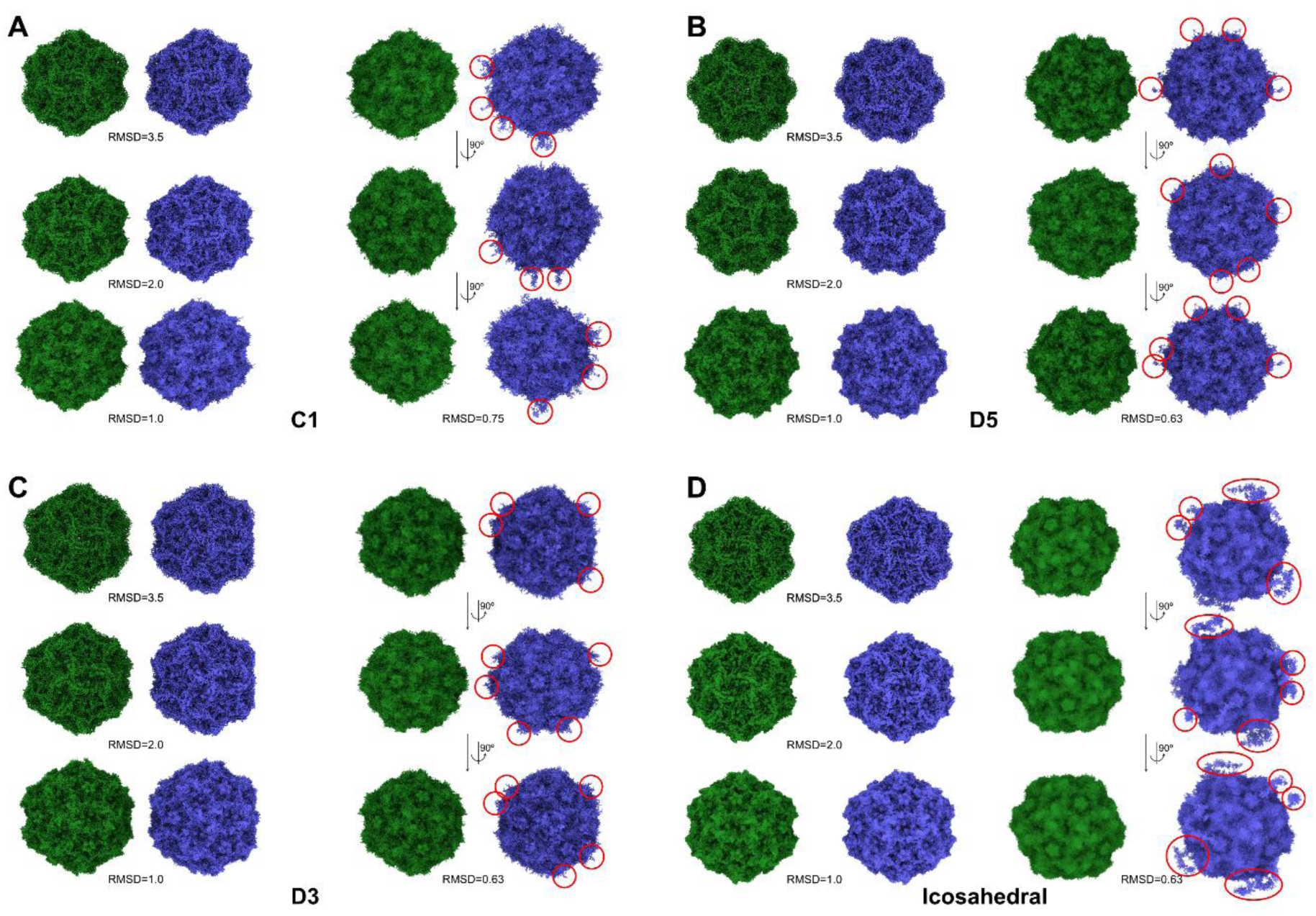
Cryo-EM reconstructions of unmodified and functionalized FPV VLPs. FPV VLP (green) and FPV VLP-peptide (blue) were analyzed. Images show four reconstructions of both FPV VLPs with different symmetries imposed: **(A)** C1, **(B)** D3, **(C)** D5 and **(D)** icosahedral. Additional densities of Coulomb potential maps on the surface of functionalized particles (red circles) are visible regardless of symmetry imposed. RMSD levels of maps contours are indicated for better readability.

### 2.6. Cell delivery of decorated FPV VLPs

FPV infects target tissues through interacting with the transferrin receptor. This characteristic of the parental virus provides an interesting potential for the derived VLP to serve as a specific cellular vector. To investigate whether our fluorescently labeled FPV VLPs do indeed specifically enter cells expressing the transferrin receptor, we tested their transducing capacity in three cell lines: CRFK (expressing feline TfR), Huh7 expressing human TfR, and HEK293T (no TfR expression). As expected, the signal from FPV VLPs could be only detected in CRFK and Huh7, but not in HEK293T providing evidence, that this transduction is receptor mediated and thus specific [**Figure 7A**].

**Figure 7.**
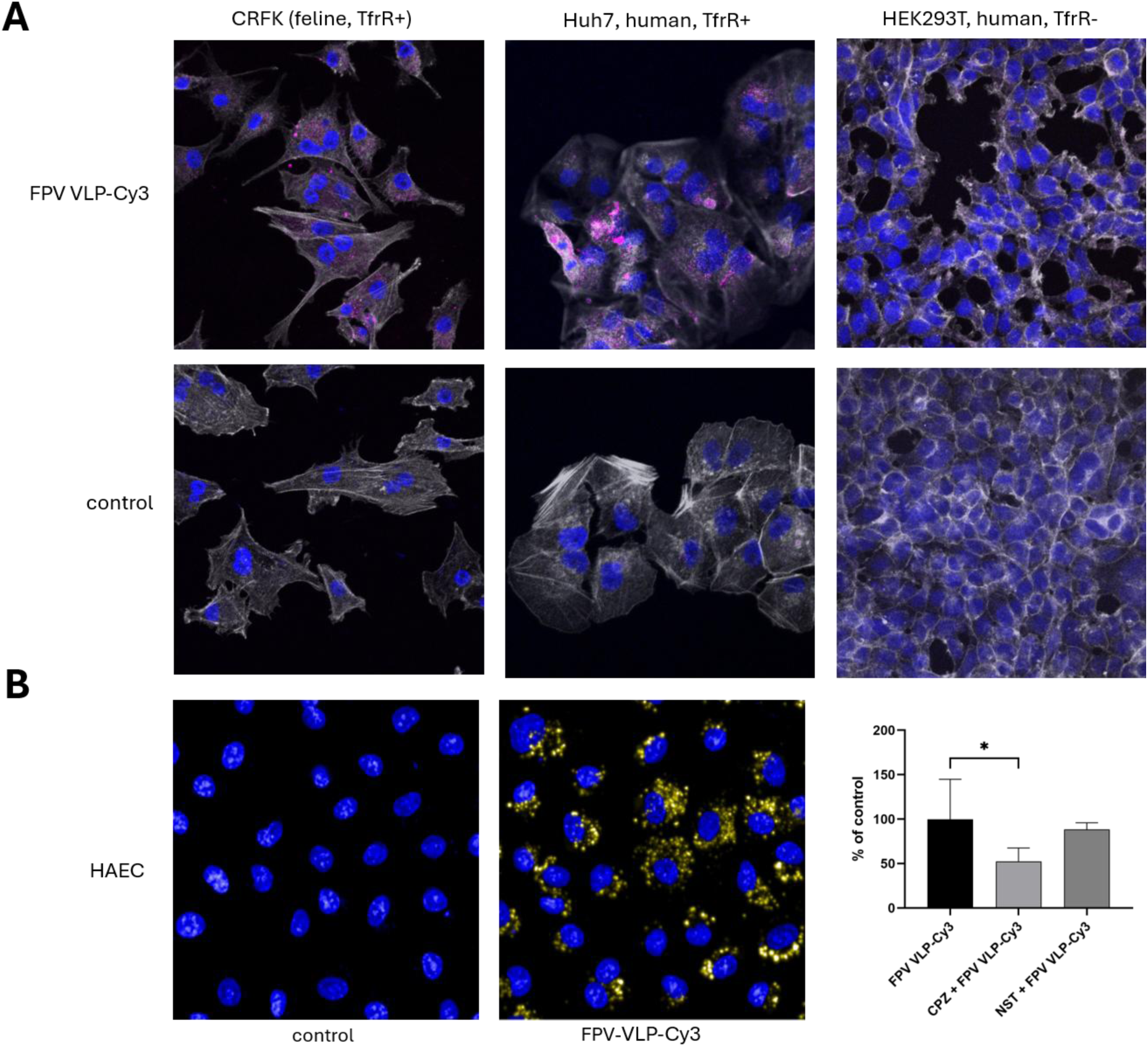
FPV VLPs enter cells expressing TfR. CRFK, Huh7 and HEK293T were incubated with FPV VLP-Cy3 (magenta), then fixed and stained to visualize actin (gray) and nuclei (blue). Preparations were imaged under confocal microscope (**A**). Human aortic endothelial cells (HAECs) were pretreated with chlorpromazine (CPZ, 10 µM) or nystatin (NST, 10 µM) and subsequently incubated with FPV VLP-Cy3 (yellow). Cells were then fixed and stained to visualize nuclei (blue). Preparations were imaged and analyzed using a CQ1 image cytometer. Images show untreated HAEC (left) and HAEC treated with FPV VLP-Cy3 only (middle). Graph shows relative quantified signal from HAEC pretreated with CPZ and NST (right). Data represent the mean ± SD of three independent experiments. Statistical analysis was performed using one-way ANOVA followed by Dunnett’s multiple comparisons test (*p < 0.05) (**B**).

Next, we confirmed that FPV VLPs retain this endocytic ability in human aortic endothelial cells (HAECs), which are considered as one on model human primary cells and which do express TfR [51]. [**Figure 7B**]. Since transferrin is internalized via receptor-mediated, clathrin-dependent endocytosis [52], we then examined whether FPV VLPs utilize this pathway in HAECs. Treatment of HAECs with chlorpromazine (CPZ), a well-established inhibitor of clathrin-mediated endocytosis, reduced FPV VLP uptake by approximately 50% [**Figure 7B**]. In contrast, treatment with nystatin, an inhibitor of caveolae-mediated endocytosis, did not affect FPV VLP uptake [**Figure 7B**]. The significant inhibition by CPZ indicates that FPV VLP internalization in HAECs is predominantly mediated by a receptor-dependent, clathrin-dependent endocytic pathway, further supporting the involvement of transferrin receptor-mediated uptake.

## 3. Discussion

In this study, we produced and functionalized virus-like particles derived from feline panleukopenia virus with the aim of creating and extensively characterizing a tissue-specific platform enabling the display and targeted delivery of various molecules. Drawing on previous fundamental studies of parvoviruses and leveraging recent advances in protein engineering, we sought to improve and further refine existing VLP-based systems.

Early works on the structural characterization of FPV VLPs relied on amplification of FPV in NLFK feline kidney–derived cells, which resulted in the production of both fully infectious viral particles and empty capsids [31]. Crystallization and subsequent structure determination of these particles provided essential information on key motifs determining host specificity [22]. Subsequent advancements in recombinant protein expression technologies enabled the production of CPV and FPV VLPs without the need to co-produce infectious viruses. Most studies report the use of the insect cell–baculovirus expression system for this purpose [35, 46, 47, 55], although assembly of CPV VLPs from VP2 protein produced in bacteria (*E. coli*) [56, 57] has also been shown to be feasible. Consequently, the purification strategies depend on the expression system used. When VLPs are assembled within producer cells and isolated as pre-assembled particles, density gradient purification (using either CsCl or sucrose gradients [35, 41, 46, 58]) is most commonly applied. In contrast, when VP2 protein is expressed in bacteria as a fusion with a purification tag (e.g., His-tag or SUMO [56, 59]) and VLP assembly occurs following tag cleavage, affinity chromatography is typically employed. In our approach, we used High Five insect cells to express codon-optimized VP2 protein, followed by purification using affinity and size-exclusion chromatography. The additional affinity purification on Ni^2+^ resin (for His-tagged VLPs) reduced the amount of contaminating proteins in samples loaded onto the SEC column, thereby improving the efficiency of this step. However, a fraction of His-tagged VLPs failed to bind to the Ni²⁺ affinity column, resulting in some material loss. Overall, both approaches (with and without affinity purification step) yielded similar amount of purified VLPs amount being approximately 20 mg of VLPs from 100 mL of insect cells culture.

Parameters such as thermal stability, aggregation propensity, and sensitivity to additives are critical for the formulation of therapeutic and preventive products, including those based on VLPs. Ongoing research aims to minimize cold-chain requirements, extend shelf life, reduce dosage, and improve immunogenicity [60–62]. In line with these objectives, we extensively investigated the stability of our FPV VLPs under elevated temperatures, across a wide range of pH conditions, and in the presence of additives commonly used to prevent aggregation. Notably, FPV VLPs demonstrated exceptional resistance to extremely low pH values (down to pH 1), which to our knowledge, represents a distinctive feature – even among VLPs - and expands their potential applicability. Buffer screening identified mildly acidic conditions (pH 4–5) as optimal for preventing thermally induced particle degradation. Furthermore, FPV VLPs exhibited good tolerance to polysorbates (Tweens) and PEG 8000, which likely additionally prevents their aggregation in long term storage and may enhance immunogenicity [63, 64]. These findings are of importance due to growing evidence that thermal stability and formulation with such additives correlate with vaccine efficacy for both preventive and therapeutic applications [65–67].

The repetitive architecture of VLPs enables high-density, spatially controlled ligand display, which is advantageous for vaccine design, targeted delivery, and biomaterials engineering. Importantly, modular VLPs offer a flexible route for introducing functional cargo without compromising particle assembly, while also enabling independent control over stoichiometry and composition. In this work, we conferred modularity of FPV VLP by targeting lysines residues in VP2 in combination with click-chemistry between the adaptor molecule (DBCO-NHS ester) and ligand of choice. Notably, functionalization of FPV VLP using this approach was more efficient than direct ligand conjugation to lysines, suggesting that DBCO-NHS ester functions not only as an adaptor, but also as a spacer molecule. This strategy was first proposed by He and Steinmetz who displayed a cancer peptide and an ODN on a plant virus derived VLP [11] and has since been adopted by other groups [9, 68]. More recently, Garcia-Truillo et al. demonstrated that this approach (DBCO-NHS ester combined with clicking azide-ligand) can be also applied to enveloped Gag-VLP [69]. Earlier studies reported VLP functionalization using click-chemistry with genetically encoded unnatural amino acids containing alkyne or azide groups, which may however negatively impact recombinant production efficiency [16]. It is also worth noting that initial click strategies relied on Cu(I)-catalyzed azide-alkyne cycloaddition [12, 15] which has since been replaced by copper-free strain-promoted azide−alkyne cycloaddition (SPAAC), eliminating concerns related to copper toxicity in biological applications.

Another critical parameter for VLP-based applications is the yield of functionalization. Here, we sought to quantify the number of molecules attached to the FPV VLP surface using gel densitometry (based on fluorescent signal), mass photometry, and RP-HPLC peak integration (for peptide conjugation). Each method has inherent limitations, which additionally depend on the type of molecule analyzed (e.g., fluorophore, peptide, ODN, protein). For instance fluorescent based quantification is biased by the degree of labeling of the molecule of interest. Moreover, SDS-PAGE densitometry is applicable only to conjugation resulting in covalent bonds (which are not disrupted by denaturation). Mass photometry, on the other hand, not only requires appropriate standard to confer linearity of the calibration curve, but also is associated with a relatively high systematic error, limiting its accuracy for small and very large molecules quantification. Finally, RP-HPLC-based estimation requires a dedicated column for each analyte. Nevertheless, the calculated average number of attached molecules was consistent with theoretical expectations [**Supplementary Table 1**]. Given that approximately 3–4 lysine residues per VP2 unit are surface-accessible, a total of 180 – 240 potential binding sites per particle can be expected. While near-complete surface occupancy is feasible for small molecules (i.e. fluorophores), steric constraints limit the number of larger ligands (peptides, ODNs, proteins) that can be accommodated. The measured amount of ODNs and peptides per VLP (~90 and ~71) falls within the range reported in the literature (20–270), depending on the VLP system, ODN length, and conjugation strategy [12]. In our study, peptide functionalization of FPV VLPs was further confirmed by cryo-EM analysis. The presence of additional densities on the VLP surface—observed regardless of the imposed symmetry during reconstruction—indicates abundant decoration. However, due to the intrinsic flexibility of the attached moieties, the structure of the modifications themselves could not be resolved. This observation is consistent with previous reports showing that peptides displayed on PP7 VLPs are not always well-resolved in cryo-EM reconstructions [70].

Having extensively characterized FPV VLPs and established their functionalization potential, we next evaluated their receptor specificity. Cross-reactivity of FPV with human transferrin receptor (TfR) has been previously reported but has not been explored in a biotechnological context. Here, we demonstrate that FPV VLPs can selectively enter human cells expressing TfR, and that this endocytosis is clathrin-mediated. This is in line with previous study describing TfR-clathrin dependent endocytosis in HAECs [51] and provides a foundation for the development of a tissue specific vector.

## 4. Conclusion

Taken together, these results establish FPV VLPs as a robust, highly stable, and modular platform with strong potential for the development of vaccines, targeted delivery vector, and system for other biotechnological applications.

## 5. Methods

### 5.1. Protein expression and purification

VP2 and His-VP2 proteins were produced in insect cells/baculovirus system. Sf9 (*Spodoptera frugiperda,* ATCC: CRL-1711) and HF (High Five, *Trichoplusia ni*, ATCC: CRL-7701) cells were cultured in ESF (Expression Systems, CA, USA) medium supplemented with 2% FBS (fetal bovine serum) (ThermoFisher Scientific), 100 µg/ml streptomycin, 100 IU/ml penicillin, 10 µg/ml gentamycin, and 0.25 µg/ml amphotericin B. The culture was maintained in a humidified incubator at 27°C. Sf9 cells were used for baculovirus (BVs) generation and amplification, while HF cells were used for recombinant protein expression. The gene encoding for VP2 protein was codon-optimized for insect and purchased as synthetic construct in pET21a (Biocat GmbH), then subcloned into pFastBac1 plasmid. The recombinant bacmid and resulting baculovirus (BV) were generated using the BAC-TO-BAC system (ThermoFisher Scientific) following manufacturer’s instructions. HF cells were then infected with recombinant BV coding for VP2 or His-VP2 proteins (at MOI = 4) and cultured for 72 hours. Proteins were extracted from insect cells by sonication, followed by centrifugation (4,500 × *g*, 20 min). The soluble fraction was then treated with Viscolase (AA Biotech, Poland) and filtered through 0.22 µm. Next, VLPs presumably present in the filtrate were separated from smaller proteins by ultrafiltration using a centrifugal device (Millipore) with MWCO = 100 kDa. The eluate was then purified on affinity (in the case of His-VP2) and SEC or directly SEC (in the case of VP2) chromatography. Namely, sample was loaded on a His-TRAP HP column (Cytiva) connected to AKTA FPLC system, followed by washing with 50 mM sodium phosphate buffer, pH 8, with 300 mM NaCl (wash buffer) and elution with a gradient of imidazole concentration (up to 500 mM) in the wash buffer. Collected fractions were analyzed for the presence of the His tag (as described in next the section), then pooled and concentrated using ultrafiltration devices (MWCO = 100 kDa, Merck-Millipore). Next, sample containing FPV VLPs (typically 10 mg/mL of total protein concentration) was separated on a Superose 6 Increase column (GE Healthcare) connected to an AKTA FPLC system, in PBS buffer.

### 5.2. VLP stability tests

Temperature resistance: VLPs (typically 0.6 mg/mL) were incubated for 60 minutes in 37°C, 50°C, 60°C, 70°C, 80°C, 95°C). Stability in pH range: aliquots of VLPs in PBS were acidified with 1 M HCl or alkalized with 1 M NaOH to reach the indicated pH value and incubated in room temperature for 60 minutes. Stability in denaturing reagents and organic solvents: VLPs were mixed with urea (reaching final concentrations: 0.5 M, 1 M, 2 M), guanidine chloride (reaching final concentrations: 0.6 M, 1.2 M, 2.4 M) SDS (reaching final V/V concentrations 50%, 25%, 12.5% for each), DMSO, DMF, (reaching final V/V concentrations 0.5%, 5%, 10%) and incubated for 120 minutes in room temperature. Stability in detergents and PEG 8000: FPV VLPs were mixed with Tween 20, Tween 80 (reaching final V/V concentrations 0.02%, 0.5%) and PEG 8000 to final W/V concentration 2% and 8% and incubated overnight in 4°C. All samples were then subjected to native PAGE as described in the separate section. Stability in buffers with different pH: VLPs were mixed with (in 1 : 4 proportion V/V) with: 50 mM Na citrate pH 4, 50 mM MES pH 5.5, 50 mM PBS pH 6.5, 50 mM HEPES pH 7.5, 50 mM TRIS pH 8.5, 50 mM TRIS pH 9.5, each containing 50 mM NaCl and incubated overnight in 4°C. Samples were then analyzed using differential scanning fluorimetry (DSF) as described in the separate section.

### 5.3. Conjugating dyes, peptides, ODNs and proteins to FPV VLPs

DBCO-NHS ester (Lumiprobe), NHS-PEG-FITC (NANOCS), fluorescein-5-maleimide (Thermo Fisher Scientific), Cy3-azide (Aldrich), Cy5-azidobenzoate-ODN (5’-TATCTAACTGCTGCGCCGCCGGGAAAATACTGTACGGTTAGA-3’ Biomers), biotin azide (PEG4 carboxamide-6-azidohexanyl biotin, Invitrogen), streptavidin-Alexa Fluor 488 conjugate (Invitrogen) were purchased as lyophilizates and dissolved in DMSO or DMF, as indicated by manufacturer. Stock solutions were kept in – 20°C and used to prepare fresh working solutions, directly before conjugating reaction.

Peptide synthesis and modification: peptide (YVCPRDFVPKGIGNKDQQIG) was synthesized at 0.1 mM scale on a Liberty Blue automated microwaved synthesizer (CEM, USA) by the Fmoc chemistry. Rink-amide resin (100-200 mesh, substitution 0.90 mmol/g) was swelled overnight with dichloromethane (DCM)/dimethylformamide (DMF) (1:1). Fmoc-deprotection was performed with 25% morpholine in DMF for 5 min at 85°C. Coupling reactions were performed as per manufacturer’s recommended protocol using DIC/Oxyma activators with a fivefold excess of Fmoc-protected amino acid derivatives for 5 min at 85°C. Double coupling was applied for all Fmoc-Arg (Pbf) residues. To optimize the cost of synthesis C-terminal 6-azidohexanoic acid was coupled to peptide-resin manually with only 2-molar excess and was carried out in the presence of HATU/HOAt/DIPEA (2:2:4 eq) for 2 hours in room temperature. Completeness of coupling was verified by Kaiser test. Cleavage from the resin and side chain deprotection were achieved by treatment with TFA/Triisopropylsilane (TIS)//water (94:3:3) for 3 hours with vigorous shaking at 30°C. The resin was filtrated and TFA solution was concentrated under a nitrogen stream. The crude peptides were precipitated by addition of cold diethyl ether, followed by centrifugation (3000 rpm, 10 min). The residue was washed with cold ether (2x) and ethyl acetate (2x). Precipitated crude peptides were dried in vacuum overnight. Crude peptides were dissolved in 8 M urea and purified on an Agilent 1260 RP-HPLC using semi-preparative C18 (10×150 mm) column (Cosmosil, Nacalai tesque). Collected peptide-containing fractions were lyophilized. Purified peptides were analyzed on an analytical C18 column (Poroshell 300SB-C18, 2.1×75mm, Agilent) in a linear gradient of 10 –80% of acetonitrile with 0.1% TFA for 30 min at flow rate 0.5 ml/min. Peak signals were detected at 220, 254 and 280 nm.

Peptide labeling with fluorescein-5-maleimide (Thermo Fisher Scientific): 0.5 mM peptide stock (in water) was diluted to 0.25 mM with 50 mM HEPES, pH 8 (final pH of the reaction mixture was 7 – 7.5) and mixed with 1 mM fluorescein-5-maleimide dissolved in DMF. The reaction was conducted in 4°C, overnight, in dark and then stopped by adding equal volume of 10 mM cysteine solution.

Conjugation/clicking reactions were conducted according to the following protocol: FPV VLP (150 nM) were mixed NHS-DBCO ester (1 mM) in the proportion to reach at least 4 x molar excess over lysines externally exposed in FPV VLP. After two hours incubation in room temperature the reaction was quenched by addition of 100 mM TRIS buffer and the unreacted NHS-DBCO ester was removed by buffer exchange to PBS, using Amicon filtering device (MWCO = 100 kDa). Next, Cy3-azide or Cy5-azidobenzoate-ODN or biotin-azide or azide-modified peptide were added to FPV VLP-DBCO (in 2 x molar excess over lysines) and the reaction was further incubated in room temperature for another 2 hours. Unreacted molecules were removed by next round of buffer exchange to PBS, using Amicon filtering device (MWCO = 100 kDa).

### 5.4. SDS-PAGE and Western blot

Protein samples were mixed with (4x concentrated) denaturing buffer containing 8% SDS and 400 mM β-mercaptoethanol and boiled for 5 min. Samples were resolved on 12% Laemmli SDS-PAGE gels. PageRulerPlus Prestained Protein Ladder (ThermoFisher Scientific) was used as protein size marker. Gels were electrotransferred to an activated PVDF membrane in 25 mM Tris, 192 mM glycine, 20% methanol buffer. The membrane was then blocked with 5% skimmed milk in Tris-buffered saline supplemented with 0.05% Tween 20, followed by 90 minutes incubation with anti-His tag antibody conjugated with horse radish peroxidase (1:20,000, Sigma). The signal was developed using a Pierce ECL Blotting Substrate (Thermo Fisher Scientific) and visualized using a gel imaging system (BioRad Chemidoc). Fluorescent signal (Cy3, FITC, Alexa488, Cy5) was detected in respective channels of Biorad Chemidoc imager.

### 5.5. Native gel electrophoresis

For non-denaturing (native) protein electrophoresis on polyacrylamide gels, precast Bis-Tris polyacrylamide 3-12% gradient gels were used (Life Technologies/ThermoFisher Scientific), following the manufacturer’s recommendations. For separating VLPs decorated with biotin-streptavidin (too large to penetrate polyacrylamide gels), 1% agarose gels were prepared in Tris-acetate buffer and electrophoresis was run under 60 V, for 90 min, in 4°C. Both polyacrylamide and agarose gels were stained with InstantBlue (Expedeon) and imaged in BioRad Chemidoc detector.

### 5.6. NanoDSF and DLS

The conformational and colloidal behaviour of VLPs was characterized on a Prometheus Panta (NanoTemper Technologies), which combines nanoDSF with dynamic light scattering (DLS) in a single measurement and thereby allows conformational and colloidal stability to be assessed in parallel. Samples of purified VLPs (with or without other molecules attached) were diluted in appropriate buffers to a final concentration of 0.05 mg/mL, filtered through 0.2 µm membrane filters (VWR) and centrifuged for 10 min at 12,045 × g. For each measurement, 10 µL of sample was loaded into high-sensitivity capillaries (PR-C006), and 2 technical replicates were run per condition.

In size analysis mode, DLS was recorded at a constant 25 °C to determine the hydrodynamic radius, size distribution, and polydispersity of the sample. In thermal mode, nanoDSF and DLS were acquired simultaneously over a temperature gradient from 25 to 95 °C at a ramp rate of 1 °C/min: the intrinsic fluorescence ratio (350/330 nm) reported the unfolding transition and apparent melting temperature, while the parallel backscatter DLS signal tracked the change in particle size and the onset of aggregation. Data were collected with PR.Panta Control software (NanoTemper Technologies, v1.4.4) and analyzed with PR.Panta Analysis software (NanoTemper Technologies, v1.4.4).

### 5.7. Transmission electron microscopy

Samples of purified VLPs (with or without other molecules attached) were diluted to 0.05 mg/mL (A280), filtered through 0.2 µm membrane filters (VWR) and centrifuged for 10 min at 12045 × g. Samples were then applied to hydrophilized carbon-coated copper grids (STEM Co.), negatively stained with 3% phosphotungestic acid (pH 7.4) and visualized using a JEOL JEM-1230 transmission electron microscope (TEM) at 80 kV. Imaging was provided as an external service by the Department of Cell Biology and Imaging from Institute of Zoology and Biomedical Research of the Jagiellonian University.

### 5.8. Cryo-EM single particle analysis experiments

Cryo-EM experiments were conducted in National cryo-EM facility at National Synchrotron Radiation Centre SOLARIS. Purified samples of FPV VLP and FPV VLP-peptide (both 150 nM) were applied on cryo-EM grids (Quantifoil, Cu 2/2, mesh 300) previously glow-discharged (70 mA, 20 s). Samples were subsequently frozen using Vitrobot IV (Thermo Fisher Scientific) with following parameters: humidity – 100 %, temperature – 4 °C, blotting force – 0, blotting time – 4 s, waiting time – 0 s, draining time – 0 s. Frozen samples were transferred to cryo-electron microscope (Titan Krios G3i) for imaging. Images of the frozen samples we collected using Gatan K3 detector equipped with BioQuantum energy filter. 11,781 movies were collected for FPV VLP-peptide and 13,489 movies were collected for non-functionalized FPV VLPs. Movies were pre-processed using motion correction (Patch Motion Correction) and contrast transfer function estimation (Patch CTF) for each of the data set [53]. As a next step, bad micrographs were discarded from the pool and the initial particle picking took place. Initially, particles were picked using blob picker from a subset of 1,000 micrographs. Picked particles were 2D-classfied (2 rounds) in order to create sensible templates for downstream template picking of the particles. There were two rounds of template picking: (i) from randomly selected 1,000 micrographs, from which particles were 2D classified and better templates were created, and (ii) from full data set using refined templates from step (i). Particles picked in step (ii) of template picking were 2D classified and final subsets of particles were selected for initial 3D reconstruction (67,788 and 157,298 particles for functionalized and non-functionalized respectively). Selected particles were further processed in following steps: *ab-initio* reconstruction (3 classes, 10,000 particles), heterogenous refinement of *ab-initio* created classes with all selected particles and finally homogenous refinement of the best class (out of 3 *ab-initio* calculated) with increasing symmetry order imposed (C1, D3, D5 and I). As a final step of coulomb potential map reconstruction, postprocessing [54] was performed (in all previously imposed symmetries). All details of the processing pipelines for both samples can be found on **Supplementary Figure 5, 6** (functionalized and non-functionalized respectively).

### 5.9. Transducing cells with fluorescently labeled FPV VLPs

Crandell feline kidney cells (Crfk, ATCC: CCL-94), Huh7 (human hepatocellular carcinoma cell line), human embryo kidney cells (HEK 293T/17, ATCC: CRL-11268) were cultured in Dulbecco’s Modified Eagle Medium (DMEM, Sigma) supplemented with 10% FBS (Sigma), 100 µg/ml streptomycin, 100 IU/ml penicillin (Sigma). Human aortic endothelial cells (HAECs, Lonza) were cultured in EGM-2 medium (Lonza) supplemented with 2% FBS. The cultures were maintained at 37 °C under 5% CO_2_. For VLP transduction, cells were grown on 15-mm glass cover slips plated into a 12-well plates (VWR) and cultured 24 – 48 hours, to reach ~60% of confluency. Next, 35 μg/well of functionalized VLPs was added and cells were incubated for 2, or 4, or 6 hours at 37°C, 5% CO_2_. After incubation, cells were washed three times for 5 min with phosphate buffered saline (PBS, EURx) and fixed with 4% paraformaldehyde solution (15 min, at room temperature). For actin staining cells were first permeabilized with 0.5% Triton-X100 in PBS (7 min, at room temperature) and then incubated with phalloidin-Alexa647 (Thermo Fisher Scientific), for 1 hour, at room temperature. After washing three times with PBS, cover slips were mounted on slides using Prolong Diamond medium with DAPI (Thermo Fisher Scientific). Fluorescent images were acquired with a Axio Observer.Z1 inverted microscope (Carl Zeiss, Jena, Germany), equipped with a LSM 880 confocal module with 40x oil immersion objective. Images were processed using ImageJ 1.47v (National Institute of Health). For studying endocytosis pathways, HAECs were seeded on 96-well plate and incubated for 24h - 48h to reach ~70% of confluency. Next, receptor-dependent, clathrin and caveolin-mediated endocytosis inhibitors, chlorpromazine (Sigma-Aldrich, St. Louis, MO, USA) and nystatin (Sigma-Aldrich, St. Louis, MO, USA) were added for 1h, following 6 μg/mL of fluorescently labeled FPV VLPs. The incubation was performed for next 24h. After incubation, cells were washed three times for 5 min with PBS (Gibco, London, UK), fixed with 4% paraformaldehyde solution (15 min, at room temperature) and stained with Hoechst 33342 (Sigma-Aldrich, St. Louis, MO, USA) to visualize nuclei. Cells were imaged using a CQ1 image cytometer (Yokogawa, Tokyo, Japan) and analysed using Columbus version 2.4.2 software (Perkin Elmer, Waltham, MA, USA).

## Supporting information

Supplemantary Data FPV VLP

## Acknowledgements

We gratefully acknowledge Polish high-performance computing infrastructure PLGrid (HPC Centre: ACK Cyfronet AGH) for providing computer facilities and support within computational grant no. PLG/2025/018508

We acknowledge Olga Woznicka for her input in TEM imaging. We are grateful to the Virogenetics Lab in MCB UJ for sharing CRFK, Huh7 and HEK293T cell lines and to Aleksandra Synowiec for her valuable contribution in transduction experiments. We thank Anna Szot for her technical help in VLP preparation and Jacek Plewka for his efforts in mass photometry data acquisition and processing.

## Data availability

Cryo-EM reconstruction of non-functionalized FPV-VLP was deposited in Electron Microscopy Data Bank under the code: EMD-58499

## References

1. Ludwig, C. and R. Wagner, Virus-like particles-universal molecular toolboxes. Curr Opin Biotechnol, 2007. 18(6): p. 537–45.

2. Lua, L.H., et al., Bioengineering virus-like particles as vaccines. Biotechnol Bioeng, 2014. 111(3): p. 425–40.

3. Nooraei, S., et al., Virus-like particles: preparation, immunogenicity and their roles as nanovaccines and drug nanocarriers. J Nanobiotechnology, 2021. 19(1): p. 59.

4. Mohsen, M.O., et al., Virus-like particles for vaccination against cancer. Wiley Interdiscip Rev Nanomed Nanobiotechnol, 2020. 12(1): p. e1579.

5. Comas-Garcia, M., M. Colunga-Saucedo, and S. Rosales-Mendoza, The Role of Virus-Like Particles in Medical Biotechnology. Mol Pharm, 2020. 17(12): p. 4407–4420.

6. Bachmann, M.F., et al., Virus-like particles: a versatile and effective vaccine platform. Expert Rev Vaccines, 2025. 24(1): p. 444–456.

7. Naskalska, A. and K. Pyrć, Virus Like Particles as Immunogens and Universal Nanocarriers. Pol J Microbiol, 2015. 64(1): p. 3–13.

8. Sitasuwan, P., et al., RGD-conjugated rod-like viral nanoparticles on 2D scaffold improve bone differentiation of mesenchymal stem cells. Front Chem, 2014. 2: p. 31.

9. Mohsen, M.O., et al., Vaccination with nanoparticles combined with micro-adjuvants protects against cancer. J Immunother Cancer, 2019. 7(1): p. 114.

10. Mohsen, M.O., et al., Targeting Mutated Plus Germline Epitopes Confers Pre-clinical Efficacy of an Instantly Formulated Cancer Nano-Vaccine. Front Immunol, 2019. 10: p. 1015.

11. Hu, H. and N.F. Steinmetz, Development of a Virus-Like Particle-Based Anti-HER2 Breast Cancer Vaccine. Cancers (Basel), 2021. 13(12).

12. Hincapie, R., et al., Preparation and Biological Properties of Oligonucleotide-Functionalized Virus-like Particles. Biomacromolecules, 2023. 24(6): p. 2766–2776.

13. Cai, H., S. Shukla, and N.F. Steinmetz, The Antitumor Efficacy of CpG Oligonucleotides is Improved by Encapsulation in Plant Virus-Like Particles. Adv Funct Mater, 2020. 30(15).

14. Laomeephol, C., et al., Surface functionalization of virus-like particles via bioorthogonal click reactions for enhanced cell-specific targeting. Int J Pharm, 2024. 660: p. 124332.

15. Banerjee, D., et al., Multivalent display and receptor-mediated endocytosis of transferrin on virus-like particles. Chembiochem, 2010. 11(9): p. 1273–9.

16. Patel, K.G. and J.R. Swartz, Surface functionalization of virus-like particles by direct conjugation using azide-alkyne click chemistry. Bioconjug Chem, 2011. 22(3): p. 376–87.

17. Brune, K.D., et al., N-Terminal Modification of Gly-His-Tagged Proteins with Azidogluconolactone. Chembiochem, 2021. 22(22): p. 3199–3207.

18. Jewett, J.C. and C.R. Bertozzi, Cu-free click cycloaddition reactions in chemical biology. Chem Soc Rev, 2010. 39(4): p. 1272–9.

19. Paradiso, P.R., S.L. Rhode, and I.I. Singer, Canine parvovirus: a biochemical and ultrastructural characterization. J Gen Virol, 1982. 62 (Pt 1): p. 113–25.

20. Parrish, C.R. and L.E. Carmichael, Antigenic structure and variation of canine parvovirus type-2, feline panleukopenia virus, and mink enteritis virus. Virology, 1983. 129(2): p. 401–14.

21. Egberink, H., et al., Vaccination and Antibody Testing in Cats. Viruses, 2022. 14(8).

22. Govindasamy, L., et al., Structures of host range-controlling regions of the capsids of canine and feline parvoviruses and mutants. J Virol, 2003. 77(22): p. 12211–21.

23. Goodman, L.B., et al., Binding site on the transferrin receptor for the parvovirus capsid and effects of altered affinity on cell uptake and infection. J Virol, 2010. 84(10): p. 4969–78.

24. Parker, J.S., et al., Canine and feline parvoviruses can use human or feline transferrin receptors to bind, enter, and infect cells. J Virol, 2001. 75(8): p. 3896–902.

25. Chapman, M.S. and M.G. Rossmann, Structure, sequence, and function correlations among parvoviruses. Virology, 1993. 194(2): p. 491–508.

26. Callaway, H.M., et al., Parvovirus Capsid Structures Required for Infection: Mutations Controlling Receptor Recognition and Protease Cleavages. J Virol, 2017. 91(2).

27. Mietzsch, M., J.J. Pénzes, and M. Agbandje-McKenna, Viruses, 2019. 11(4).

28. López-Astacio, R.A., et al., The Structures and Functions of Parvovirus Capsids and Missing Pieces: the Viral DNA and Its Packaging, Asymmetrical Features, Nonprotein Components, and Receptor or Antibody Binding and Interactions. J Virol, 2023. 97(7): p. e0016123.

29. Parrish, C.R., Mapping specific functions in the capsid structure of canine parvovirus and feline panleukopenia virus using infectious plasmid clones. Virology, 1991. 183(1): p. 195–205.

30. Chang, S.F., J.Y. Sgro, and C.R. Parrish, Multiple amino acids in the capsid structure of canine parvovirus coordinately determine the canine host range and specific antigenic and hemagglutination properties. J Virol, 1992. 66(12): p. 6858–67.

31. Agbandje, M., et al., Structure determination of feline panleukopenia virus empty particles. Proteins, 1993. 16(2): p. 155–71.

32. Wu, H. and M.G. Rossmann, The canine parvovirus empty capsid structure. J Mol Biol, 1993. 233(2): p. 231–44.

33. Tsao, J., et al., Structure determination of monoclinic canine parvovirus. Acta Crystallogr B, 1992. 48 (Pt 1): p. 75–88.

34. Casal, J.I., Use of parvovirus-like particles for vaccination and induction of multiple immune responses. Biotechnol Appl Biochem, 1999. 29 (Pt 2): p. 141–50.

35. Rueda, P., et al., Engineering parvovirus-like particles for the induction of B-cell, CD4(+) and CTL responses. Vaccine, 1999. 18(3-4): p. 325–32.

36. Rueda, P., et al., Minor displacements in the insertion site provoke major differences in the induction of antibody responses by chimeric parvovirus-like particles. Virology, 1999. 263(1): p. 89–99.

37. Feng, H., et al., Recombinant canine parvovirus-like particles express foreign epitopes in silkworm pupae. Vet Microbiol, 2011. 154(1-2): p. 49–57.

38. Wang, C., et al., Novel chimeric virus-like particles vaccine displaying MERS-CoV receptor-binding domain induce specific humoral and cellular immune response in mice. Antiviral Res, 2017. 140: p. 55–61.

39. Zhao, S., et al., Development and efficacy evaluation of remodeled canine parvovirus-like particles displaying major antigenic epitopes of a giant panda derived canine distemper virus. Front Microbiol, 2023. 14: p. 1117135.

40. Muthuraman, K.R., et al., Tetravalent Virus-like Particles Engineered To Display Envelope Domain IIIs of Four Dengue Serotypes in Silkworm as Vaccine Candidates. Biomacromolecules, 2025. 26(3): p. 2003–2013.

41. Singh, P., et al., Canine parvovirus-like particles, a novel nanomaterial for tumor targeting. J Nanobiotechnology, 2006. 4: p. 2.

42. Singh, P., Tumor targeting using canine parvovirus nanoparticles. Curr Top Microbiol Immunol, 2009. 327: p. 123–41.

43. Jiao, C., et al., Construction and Immunogenicity of Virus-Like Particles of Feline Parvovirus from the Tiger. Viruses, 2020. 12(3).

44. Wang, T., et al., Virus-like Particle Vaccine for Feline Panleukopenia: Immunogenicity and Protective Efficacy in Cats. Vaccines (Basel), 2025. 13(7).

45. Feng, E., et al., Generation and Immunogenicity of Virus-like Particles Based on the Capsid Protein of a Chinese Epidemic Strain of Feline Panleukopenia Virus. Vet Sci, 2025. 12(5).

46. Gilbert, L., et al., Assembly of fluorescent chimeric virus-like particles of canine parvovirus in insect cells. Biochem Biophys Res Commun, 2004. 313(4): p. 878–87.

47. Gilbert, L., et al., Truncated forms of viral VP2 proteins fused to EGFP assemble into fluorescent parvovirus-like particles. J Nanobiotechnology, 2006. 4: p. 13.

48. Biela, A.P., et al., Programmable polymorphism of a virus-like particle. Commun Mater, 2022. 3: p. 7.

49. Malay, A.D., et al., An ultra-stable gold-coordinated protein cage displaying reversible assembly. Nature, 2019. 569(7756): p. 438–442.

50. Simpson, A.A., et al., Host range and variability of calcium binding by surface loops in the capsids of canine and feline parvoviruses. J Mol Biol, 2000. 300(3): p. 597–610.

51. Lu, S. and L. Yu, Salvianic acid A ameliorates ox-LDL-induced ferroptosis in vascular endothelial cells by regulating TFRC. Medicine (Baltimore), 2025. 104(44): p. e45542.

52. Traub, L.M., Tickets to ride: selecting cargo for clathrin-regulated internalization. Nat Rev Mol Cell Biol, 2009. 10(9): p. 583–96.

53. Punjani, A., et al., cryoSPARC: algorithms for rapid unsupervised cryo-EM structure determination. Nat Methods, 2017. 14(3): p. 290–296.

54. Zivanov, J., T. Nakane, and S.H.W. Scheres, A Bayesian approach to beam-induced motion correction in cryo-EM single-particle analysis. IUCrJ, 2019. 6(Pt 1): p. 5–17.

55. López de Turiso, J.A., et al., Recombinant vaccine for canine parvovirus in dogs. J Virol, 1992. 66(5): p. 2748–53.

56. Xu, J., et al., Self-assembly of virus-like particles of canine parvovirus capsid protein expressed from Escherichia coli and application as virus-like particle vaccine. Appl Microbiol Biotechnol, 2014. 98(8): p. 3529–38.

57. Nan, L., et al., Trigger factor assisted self-assembly of canine parvovirus VP2 protein into virus-like particles in Escherichia coli with high immunogenicity. Virol J, 2018. 15(1): p. 103.

58. Hashemzadeh, M.S. and N. Gharari, Biosynthesis of a VLP-type nanocarrier specific to cancer cells using the BEVS expression system for targeted drug delivery. J Genet Eng Biotechnol, 2023. 21(1): p. 20.

59. Yan, D., et al., Quantum Dots Encapsulated with Canine Parvovirus-Like Particles Improving the Cellular Targeted Labeling. PLoS One, 2015. 10(9): p. e0138883.

60. Mellid-Carballal, R., et al., Viral protein nanoparticles (Part 1): Pharmaceutical characteristics. Eur J Pharm Sci, 2023. 187: p. 106460.

61. Le, D.T. and K.M. Müller, In Vitro Assembly of Virus-Like Particles and Their Applications. Life (Basel), 2021. 11(4).

62. Wu, Z., et al., Optimizing the Stability of Viral Nanoparticles: Engineering Strategies, Applications, and the Emerging Concept of the Virophore. J Am Chem Soc, 2026. 148(2): p. 2081–2095.

63. Shi, S., et al., Vaccine adjuvants: Understanding the structure and mechanism of adjuvanticity. Vaccine, 2019. 37(24): p. 3167–3178.

64. Tenchov, R., J.M. Sasso, and Q.A. Zhou, PEGylated Lipid Nanoparticle Formulations: Immunological Safety and Efficiency Perspective. Bioconjug Chem, 2023. 34(6): p. 941–960.

65. Aguado-Garcia, D., et al., Evaluation of the Thermal Stability of a Vaccine Prototype Based on Virus-like Particle Formulated HIV-1 Envelope. Vaccines (Basel), 2022. 10(4).

66. Yuan, H., et al., The pH stability of foot-and-mouth disease virus. Virol J, 2017. 14(1): p. 233.

67. Lin, Y., et al., Nanovaccines empowering CD8. Theranostics, 2025. 15(7): p. 3098–3121.

68. Wang, Y., et al., Development and characterization of a human papillomavirus-based nanoparticle carrier for heterologous vaccine antigens. Sci Rep, 2025. 15(1): p. 32927.

69. García-Trujillo, M., et al., Gag HIV-1 Virus-like Particles and Extracellular Vesicles Functionalization with Spike Epitopes of SARS-CoV-2 Using a Copper-Free Click Chemistry Approach. Bioconjug Chem, 2025. 36(3): p. 486–499.

70. Zhao, L., et al., Engineering the PP7 Virus Capsid as a Peptide Display Platform. ACS Nano, 2019. 13(4): p. 4443–4454.

