## Supplementary material for "Modular Surface Engineering of Feline Parvovirus Virus-Like Particles for Antigen Presentation and Receptor-Mediated Cell Targeting": Supplemantary Data FPV VLP

Supplementary data

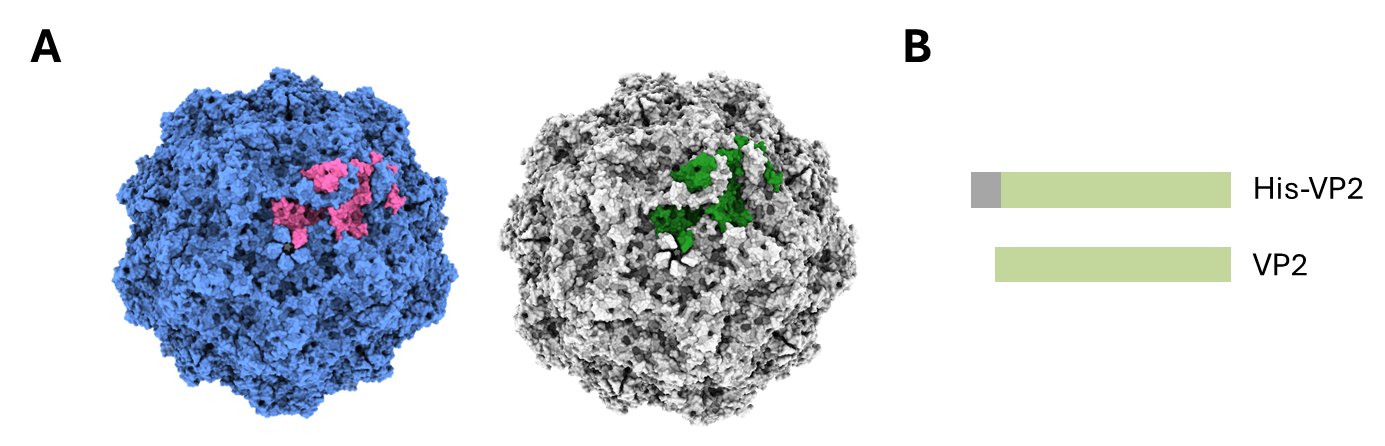

**Supplementary Figure 1. Designing FPV VLP.** CPV-derived (left) and FPV-derived (right) VLPs exhibit high structural similarity; VP2 monomer was colored in pink in CPV and green in FPV. Images were created in ChimeraX from 1C8D (CPV) and 1C8E (FPV) (**A**). FPV VLPs described in this work were assembled either from VP2 or His-VP2 proteins. His-tag (6xHis) was appended to N-terminus of VP2. (**B**).

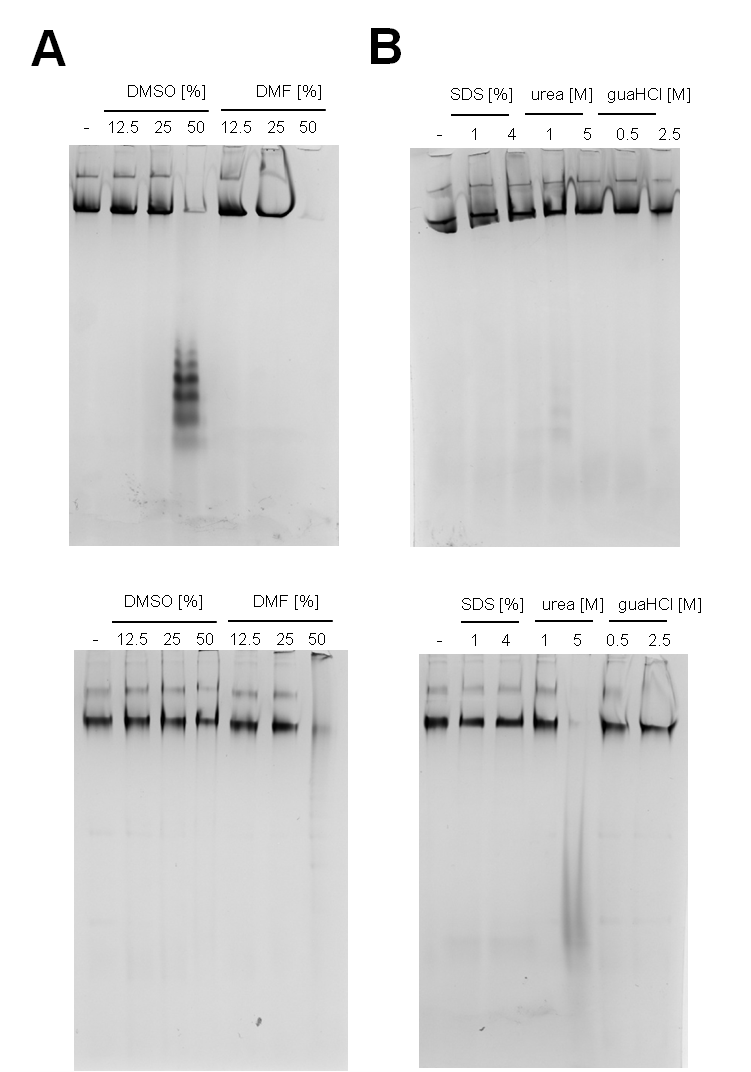

**Supplementary Figure 2.** **Stability of FPV VLPs in organic solvents and denaturing agents**. Stability of FPV VLPs (upper) and His-FPV VLPs (lower) in DMSO and DMF (**A**); sodium dodecyl sulfate (SDS), urea and guanidine hydrochloride (guaHCl) (**B**) was assessed by native PAGE, as indicated.

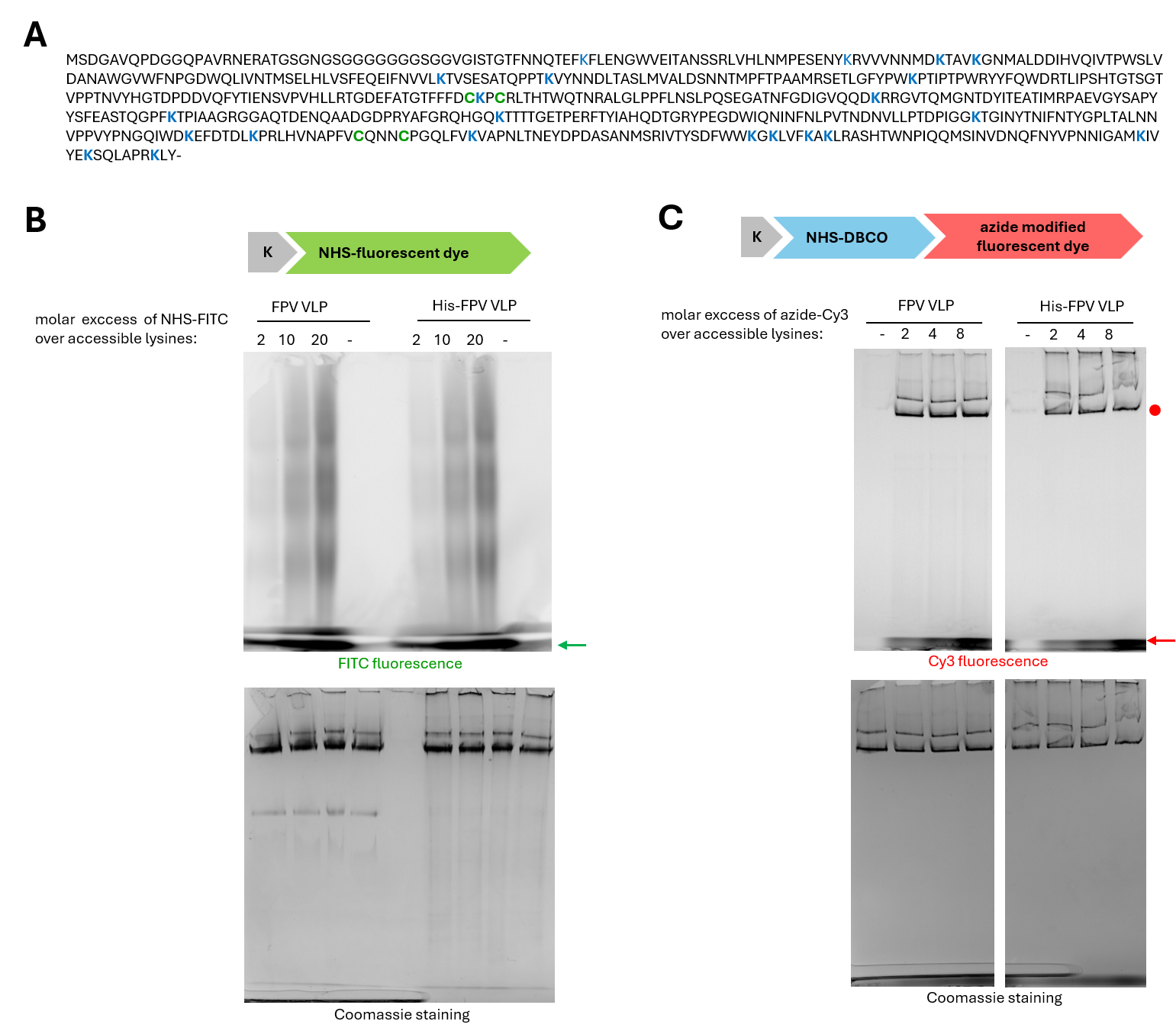

**Supplementary Figure 3. Conjugation of fluorophore to FPV and His-FPV VLPs.** Amino acids sequence of VP2 protein used for FPV VLP formation in this study. Lysines (K) are highlighted in blue and cysteines (C) – in green (**A**). Direct conjugation of FITC-PEG-NHS to lysines within FPV VLP was inefficient (**B**), whereas combining conjugation of NHS-DBCO with subsequent click reaction of azide-modified Cy3 resulted in efficient FPV and His-FPV VLP labeling (**C**). Native-PAGE gels show separation of FPV and His-FPV VLPs after conjugation reactions. Numbers indicate molar excess of fluorescent dyes over theoretically accessible lysines (here assumed as 210 – median value within 180 -240 range) on VLP surface. Gels were imaged in ChemiDoc fluorescence device (B, C upper panels) to detect FITC and Cy3 signals respectively, and then stained to detect protein bands (B, C upper panels). Arrows indicate the excess of unreacted fluorophore; red dot indicates fluorescent signal of Cy3-labeled FPV and His-FPV VLPs.

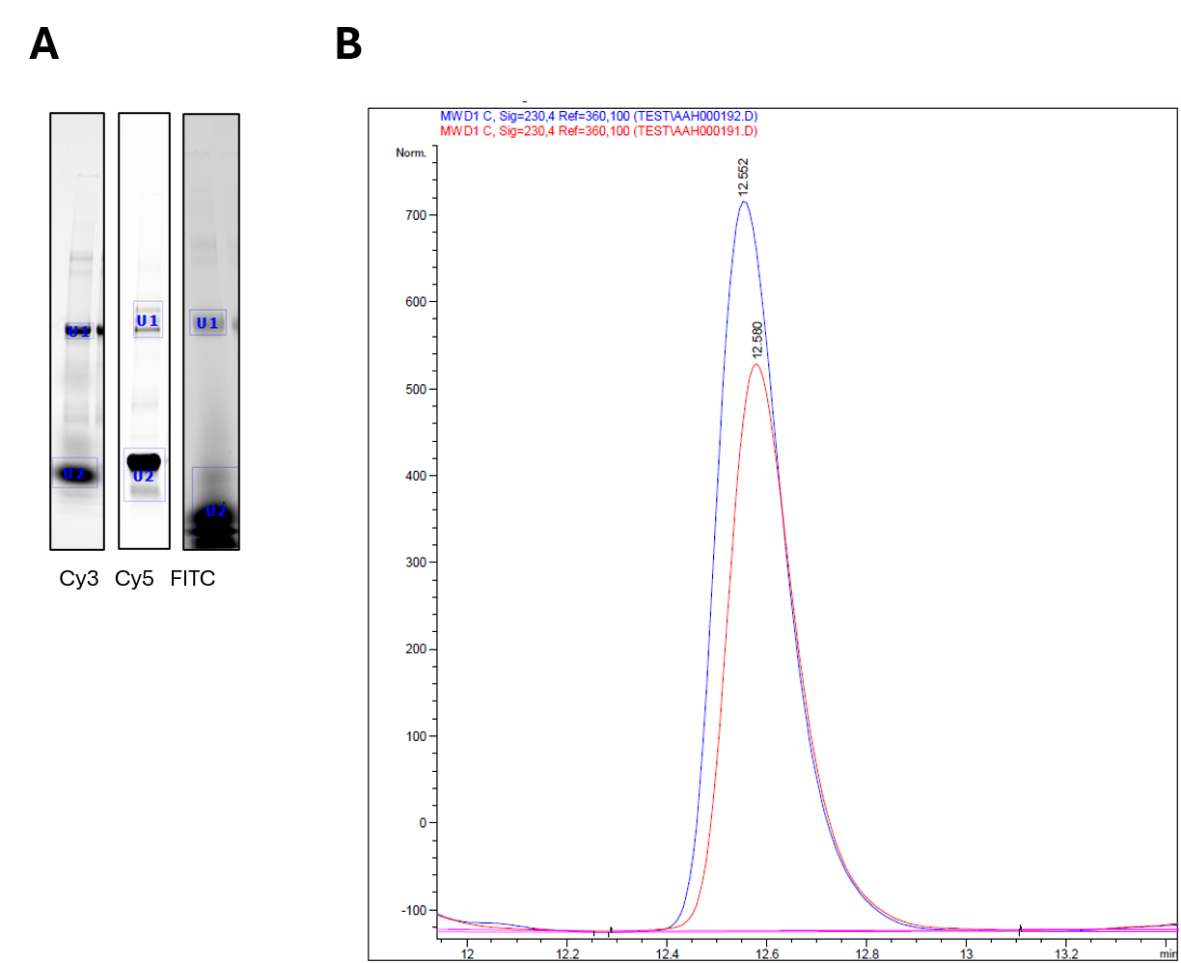

**Supplementary Figure 4. Estimation of the yield of FPV VLP decoration.** Approximate number of molecules attached to FPV VLP was determined from quantification of fluorescent signal intensity after SDS-PAGE sample separation (**A**) and reverse phase - high performance liquid chromatography (RP-HPLC) (**B**).

**A**. Intensity of the fluorescent signal measured in gels shown in Figure 4 of the main manuscript and here as inverted signal images was quantified using Imagelab software (Biorad). Fractions of reacted molecules (ligands) were calculated using the formula:

(1) Intensity_U1_/( Intensity_U1_ + Intensity_U2_) *100 = fraction of ligands attached to FPV VLP

Amount of ligands attached to FPV VLP was calculated using the formula:

(2) amount of ligands attached to FPV VLP = (amount of ligands in the reaction * fraction of ligands attached to FPV VLP)/ amount of FPV VLP in the reaction

**B**. Determination of yield of peptide conjugation to FPV VLP was additionally based on calculation of initial and final (after conjugation reaction) peptide concentration measured by RP-HPLC analysis. Decrease of peptide amount remanent (unbound) in the sample after mixing (FPV VLP + peptide) was due to the peptide attachment to FPV VLP. Samples of peptide before (initial, blue line) and after conjugation to FPV VLP (final, red line) were analyzed by RP HPLC on Agilent 1260 Infinity instrument on Agilent Poroshell C18 column (5 µm 300SB, 2.1x75 mm). The initial and final peptide concentrations were calculated by measuring the integrated peak areas detected at 210 and 280 nm using Agilent ChemStation software. Fraction of reacted peptide was calculated using formula (3) and the amount of peptide attached to FPV VLP was calculated using formula (2).

(3) (AUC_i_- AUC_f_)/AUC_i_ )*100 = fraction of peptide attached to FPV VLP

| **Sample /Method** | **SDS-PAGE + fluorescent detection** | **RP-HPLC** | **Average** |
| --- | --- | --- | --- |
| FPV VLP-Cy3 | 136 | nd | 136 |
| FPV VLP-ODN-Cy5 | 90 | nd | 90 |
| FPV VLP-peptide-FITC | 39* | 54 | 71* |

**Supplementary Table 1** Calculated amount of molecules conjugated to FPV VLP. *assuming degree of ODN or peptide labeling 50%, nd-not determined.

| **sample** | **Diameter [nm]** | **Sigma [nm]** |
| --- | --- | --- |
| FPV VLP | 28.46 | 0.15 |
| FPV VLP control (DBCO only) | 33.72 | 0.14 |
| FPV VLP-Cy3 | 31.72 | - |
| FPV VLP-peptide-FITC | 42.46 | 3.12 |
| FPV VLP-ODN-Cy5 | 36.32 | 0.02 |
| FPV VLP-biot-strept-Alexa488 | 42.22 | 0.62 |

**Supplementary Table 2.** Hydrodynamic diameters of decorated FPV VLPs measured in DLS.

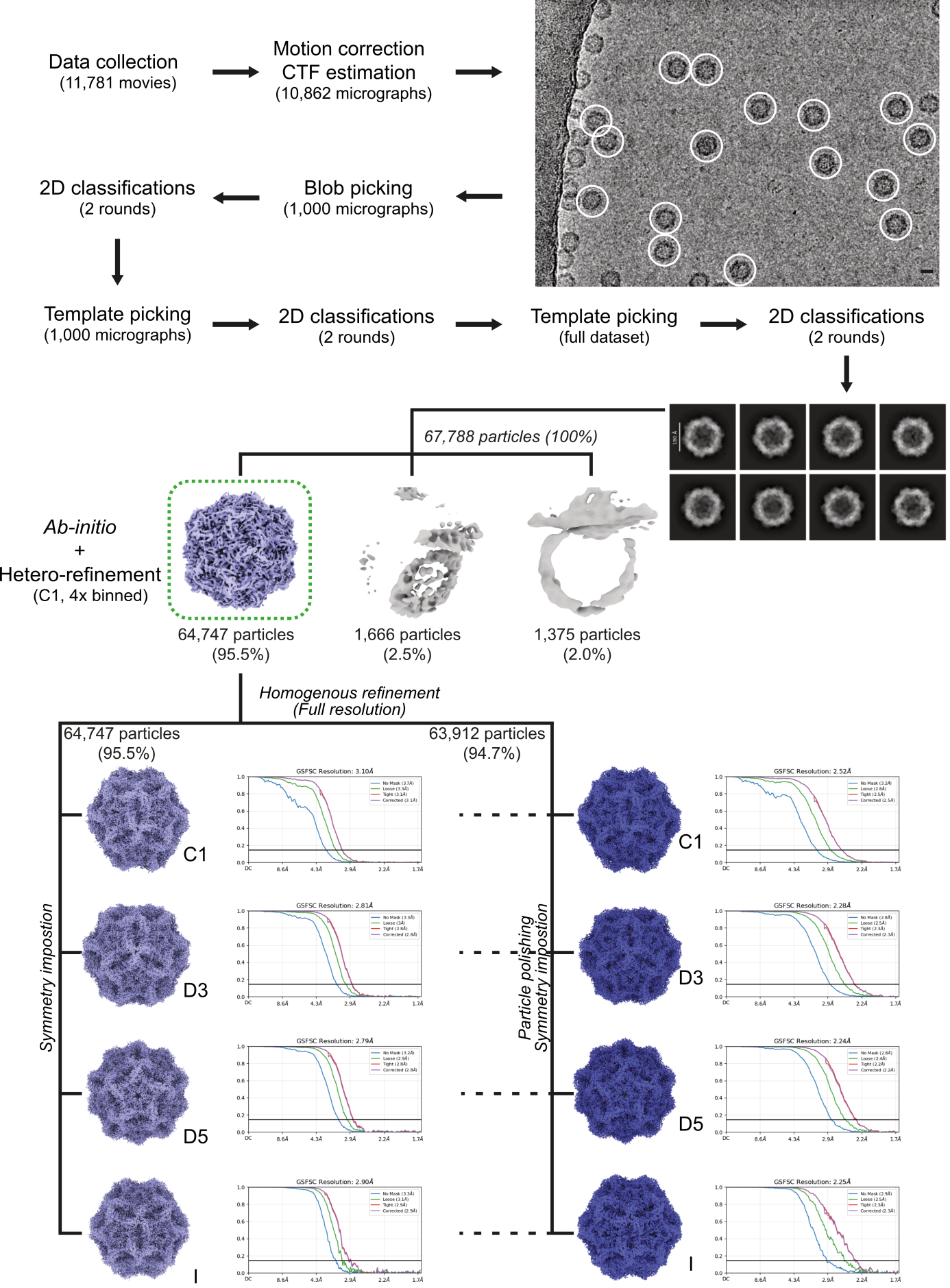

**Supplementary Figure 5.** Schematic processing pipeline of the functionalized FPV-VLP showing consecutive steps of reconstruction, starting from the collected data, and finishing on final 3D volume reconstruction with different symmetries imposed (from C1 to Icosahedral).

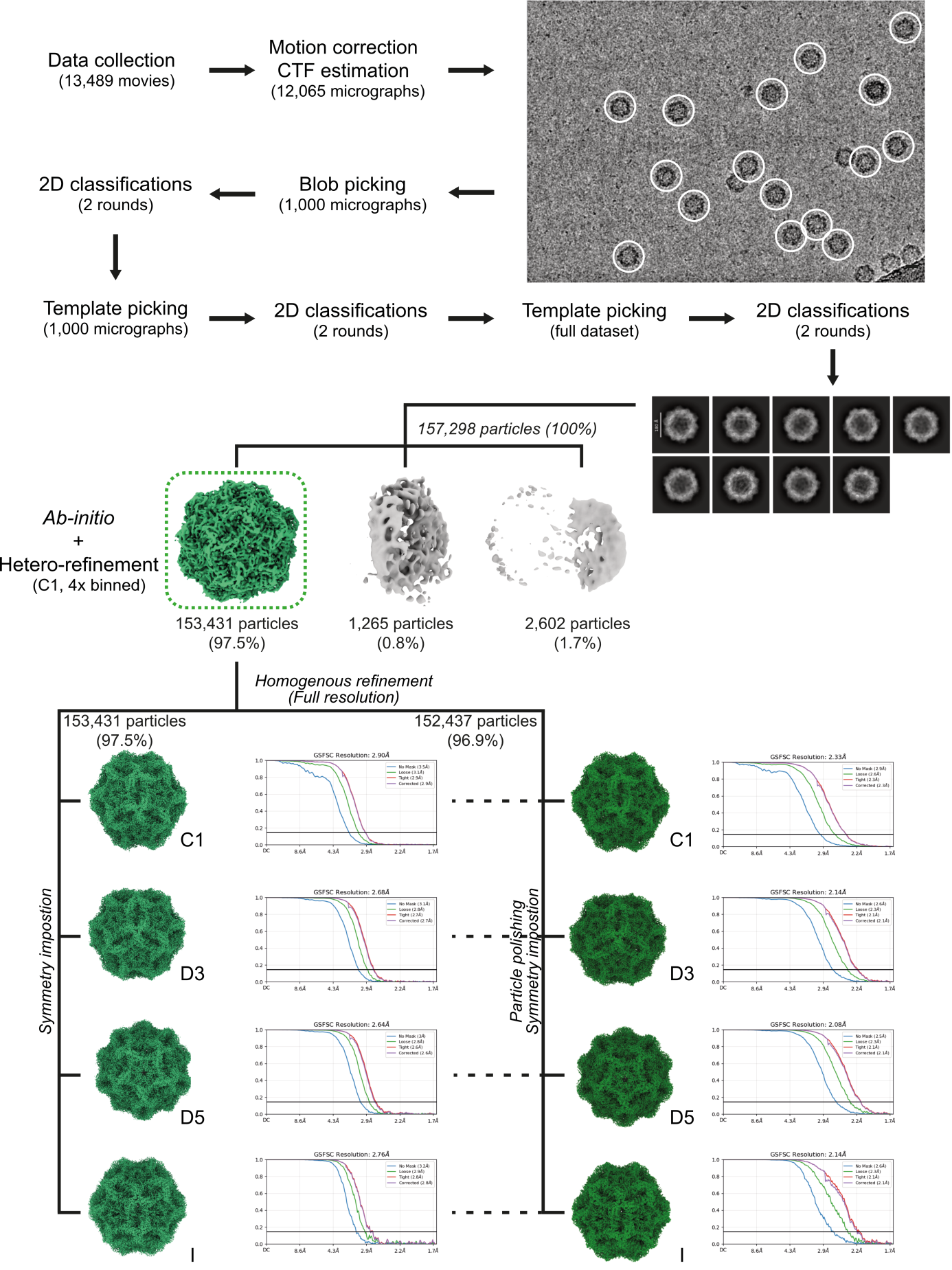

**Supplementary Figure 6.** Processing pipeline of the non-functionalized FPV-VLP showing consecutive steps of reconstruction, starting from the collected data, and finishing on final 3D volume reconstruction with different symmetries imposed (from C1 to Icosahedral).
